# Impaired Control Symptoms in Alcohol Use Disorder Show Convergent Associations Across Behavioral and Neural Markers of Inhibitory Control and Drinking Behavior

**DOI:** 10.64898/2026.10.05.756706

**Authors:** Justin Böhmer, Pablo Reinhardt, Marco Bottino, Christoph Bärtl, Fabian Arntz, Malin K. Hildebrandt, Hao Chen, Louis Thill, Johannes Petzold, Kerstin Ritter, Michael Rapp, Falk Kiefer, Patrick Bach, Bernd Lenz, Andreas Heinz, Gianna Spitta, Annika Rosenthal, Tanja Endrass, Michael N. Smolka, Rainer Spanagel, Gina-Isabelle Henze, ReCoDe-Consortium, Michael Marxen, Henrik Walter

## Abstract

**Background:** Deficits in inhibitory control are considered a central feature of alcohol use disorder (AUD), yet behavioral and neuroimaging findings have been inconsistent. One potential explanation is the marked heterogeneity of AUD, which encompasses multiple symptom dimensions that may differ in their underlying neurocognitive mechanisms. In this study, we investigated whether behavioral and neural markers of inhibitory control are associated with specific DSM-5 AUD symptom domains.

**Methods:** A total of 589 participants (432 with non-treatment seeking AUD, 157 controls) completed a Stop-Signal Task (SST) during fMRI. Associations between DSM-5 symptom domains (impaired control over alcohol use, risky use, social problems, physical dependence) and SST performance, inhibition-related brain activation, and alcohol consumption were examined. Real-life drinking behavior was additionally assessed over 12 months using ecological momentary assessment (EMA).

**Results:** Greater impaired control symptom severity was associated with poorer performance on go trials and diminished post-error slowing during SST. At the neural level, higher impaired control symptom severity was associated with reduced activation across fronto-insular, parietal, cingulate, supplementary motor, and cerebellar regions. In contrast, other symptom domains showed little to no associations with inhibition-related behavioral or neural measures. Higher impaired control symptom severity was associated with greater alcohol consumption across questionnaire-based and EMA-based assessments.

**Conclusions:** In this large cohort, altered behavioral performance and recruitment of inhibition-related brain regions in the SST was selectively linked to impaired control symptoms. These findings highlight the importance of symptom-based approaches for understanding heterogeneity in addiction and may inform the development of more targeted interventions.

## Introduction

Alcohol use disorder (AUD) ranks among the most prevalent and devastating psychiatric conditions worldwide, affecting approximately 7% of the global adult population^1^ and substantially contributing to morbidity and mortality^2^. Clinically, AUD is characterized by a persistent pattern of alcohol consumption despite adverse consequences, reflecting loss of control over drinking behavior. A key cognitive mechanism underlying this loss of control is inhibitory control, the ability to withhold automatic or prepotent responses in favor of behavior aligned with long-term goals^3–5^. Impaired inhibitory control has been identified as a key vulnerability factor for the development and maintenance of addictive behaviors,^6,7^ and is associated with higher alcohol consumption, increased craving, and greater relapse risk in AUD.^8–10^

Inhibitory control is typically measured using paradigms requiring inhibition of a dominant motor response, such as the Stop-Signal Task (SST) or the Go/No-Go task.^11,12^ Behavioral studies in AUD frequently report poorer inhibitory performance, reflected in slower stopping latencies, more commission errors, and reduced adaptive post-error adjustments.^13–16^ However, a substantial number of studies have failed to detect such deficits,^17–23^ raising questions about their robustness.

Neuroimaging research similarly points to alterations in neural systems supporting response inhibition in AUD, including reduced activation in regions such as the inferior frontal gyrus, dorsal anterior cingulate cortex, anterior insula, and supramarginal gyrus during inhibitory demands.^18,20,24–28^ Yet, meta-analytic evidence indicates considerable heterogeneity across studies examining inhibitory-related neural activation in substance use disorders, including AUD.^19,29,30^ Collectively, these inconsistent behavioral and neural findings suggest that inhibitory dysfunction may not be uniformly expressed across individuals with AUD.

One factor that may contribute to such variability is the marked heterogeneity of symptoms in AUD as currently defined. Based on diagnostic criteria of the fifth edition of the Diagnostic and Statistical Manual of Mental Disorders (DSM-5),^31^ an AUD diagnosis can be met through more than 2,000 unique symptom combinations. These symptoms cluster into four domains (**Fig. S1**): *impaired control over substance use* (e.g., drinking more than intended), *social problems* (e.g., failure to fulfill role obligations), *risky use* (e.g., recurrent use in hazardous situations), and *physical dependence* (e.g., withdrawal symptoms).^31^ Importantly, evidence indicates that different symptom profiles in AUD are linked to at least partially distinct cognitive and neurobiological processes.^32–35^ Treating AUD as a unitary category therefore aggregates individuals with diverse symptom presentations, potentially obscuring symptom-specific associations with inhibitory dysfunction and contributing to inconsistent empirical findings. From a theoretical standpoint, inhibitory control deficits may be particularly relevant to symptoms that directly reflect difficulties in regulating alcohol consumption, such as drinking more or for longer than intended and a persistent desire or unsuccessful efforts to cut down or control alcohol use. Those criteria fall within the DSM-5 *impaired control* domain and arguably show a more direct conceptual link to inhibitory control than other AUD symptoms, such as *social problems*, *risky use*, or *physical dependence*. However, to our knowledge, no study to date has tested whether inhibitory dysfunction varies systematically across DSM-5 symptom domains.

The present study addresses this gap by investigating behavioral and neural markers of response inhibition in a large sample spanning the full spectrum of AUD symptom severity, including non-treatment seeking individuals meeting DSM-5 criteria for AUD and controls. Using the SST as a validated measure of inhibitory control,^36–38^ we test whether behavioral performance and corresponding neural activation patterns are selectively associated with *impaired control* symptoms, but unrelated to symptoms reflecting *social problems*, *risky use*, or *physical dependence*. We further examine whether these symptom domains differentially relate to alcohol consumption assessed via questionnaires and prospectively in daily life via ecological momentary assessments (EMA). By linking inhibition-related behavior, brain activation, and alcohol use to distinct AUD symptom domains, this work aims to clarify the heterogeneity underlying inhibitory control deficits in AUD.

## Methods and Materials

### Participants

Participants were enrolled as part of a large German collaborative research centre on losing and regaining control over drug intake (TRR265 ReCoDe).^39,40^ Individuals aged 16–65 years were recruited across three sites (Berlin, Dresden, Mannheim) to investigate behavioral, cognitive, and neurobiological mechanisms underlying AUD, as well as its developmental trajectories and potential interventions. The present study evaluates the Stop-Signal Task-based fMRI data from this cohort, which were available for 802 participants. Individuals were not included in the study if they reported acute withdrawal symptoms, medically supervised withdrawal, current intention to seek treatment, medical history of bipolar disorder, psychotic or schizophrenia-spectrum disorder, or any substance use disorder other than AUD, tobacco use disorder, or cannabis use disorder, as well as a history of severe head injury, other central nervous system (CNS) disorders, the use of medications affecting the CNS, and contraindications to MRI scanning. Moreover, altogether 213 participants (26.56%) were excluded, either due to insufficient fMRI data quality (n = 60), incomplete task data (n = 67), or due to not meeting predefined behavioral performance criteria in the Stop-Signal Task (n = 86), resulting in a final sample of 589 participants (40.75% females) with a mean age of 35.16 years (*SD* = 12.19). Of these, 432 fulfilled two or more DSM-5 criteria and were classified as AUD subjects. All participants provided written informed consent prior to study participation. The study was conducted in accordance with the Declaration of Helsinki and approved by the medical ethics committees at each study site (EA1/212/18 [Berlin], EK 459112018 [Dresden], 2018-621N-MA [Mannheim]).

### Diagnostic and psychometric assessment

Diagnostic assessment was conducted using the Structured Clinical Interview for DSM-5 (SCID-5). For each participant, the total number of AUD criteria and the number of criteria for each symptom domain (impaired control, social problems, risky use, and physical dependence) were recorded.^31^ At baseline, alcohol use was assessed using standardized self-report questionnaires capturing the quantity and frequency of alcohol consumption during the preceding three months. For each reported alcoholic beverage, the corresponding amount of pure alcohol (in grams) was calculated. Based on these data, average daily alcohol intake, reflecting the mean amount of alcohol consumed on an average drinking day, as well as drinking frequency, defined as the percentage of days on which participants consumed at least one alcoholic beverage, and binge-drinking frequency, defined as the percentage of days involving the consumption of five or more standard drinks, was calculated.

### Ecological Momentary Assessment (EMA) of alcohol use behavior

A subset of participants meeting at least two DSM-5 criteria for AUD was followed up for 365 days using ecological momentary assessment (EMA). Using a smartphone application, participants were asked every other day to report on the quantity and type of alcoholic beverages consumed on each of the preceding two days from a list of alcohol-containing drinks. Participants were prompted to complete the assessment at 12 pm, with the option to postpone until 8 pm. Based on these reports, daily alcohol intake (in grams of pure alcohol) was calculated for each day. In addition, each day was classified as a drinking day (yes/no) or a binge-drinking day (yes/no) based on the reported amounts of alcohol intake. See Deeken et al.^41^ for further details on the EMA procedure. Participants were included in EMA-based analyses if they provided at least 90 days of valid drinking data during the 365-day follow-up period. Altogether, 45,173 entries from 389 participants with AUD were analyzed. The mean follow-up duration, defined as the interval between the first and last completed EMA assessment, was 317.62 days (SD = 76.30). Compliance rate was 80.89% (SD = 18.97) of all prompts.

### fMRI data acquisition and preprocessing

MRI data were collected on 3T Siemens Prisma/Prisma Fit and Trio scanners in Berlin, Dresden, and Mannheim, with a 64-channel coil in Berlin and Mannheim, and a 32-channel coil in Dresden. Extensive information on data acquisition parameters and preprocessing is provided in the Supplementary Information (see Extended Methods).

### The Stop-Signal Task (SST)

During fMRI, participants underwent an established version of the Stop-Signal Task (SST), in which they were instructed to respond to the direction of an arrow pointing either left or right (i.e., go signal) by pressing a button with their left or right index finger, consistent with the direction of the arrow. Infrequently, an upward-pointing arrow (i.e., stop signal) was presented after the go signal, in which subjects were instructed to withhold their already started motor response. The interval between the go and stop signal (i.e., stop-signal delay [SSD]) was continuously adjusted according to each participant’s performance. This adaptive tracking procedure gradually stabilized each individual’s probability of successful inhibition (successful stop trials / all stop trials) at approximately 50%.^42^ Further information on the timing of stimuli and number of trials is presented in the Supplementary Information (see Extended Methods). Based on published recommendations,^36^ we applied lenient criteria for identification of outlier subjects (accuracy on stop trials [inhibition rate] outside 25–75%, go rate < 60%, accuracy on go trials < 90%, Stop-Signal Reaction Time (SSRT) < 50 ms), resulting in the exclusion of 67 participants with AUD and 19 controls.

### Behavioral measures of SST performance

Several behavioral performance parameters were extracted from the SST. First, we investigated the SSRT. Specifically, SSRT was estimated based on the independent race model, which conceptualizes stopping as the latency required to interrupt an ongoing response. In this regard, all Go RTs were sorted from fastest to slowest, and the Go RT corresponding to the individual probability of responding on stop trials was identified. From this value, the participant’s mean SSD was subtracted, yielding the SSRT, with higher values indicating worse inhibition performance.^43^ Second, we evaluated the Balanced Integration Score (BIS). BIS is a composite index of overall task performance on go trials that combines speed and accuracy into a single measure by subtracting z-standardized RTs from z-standardized accuracies,^44,45^ with higher scores indicating better task performance. Third, we explored behavioral adaptation after committing errors by investigating post-error slowing (PES). PES reflects the tendency to slow down responses after committing an error in order to increase the probability of a correct response at the expense of response speed.^46^ PES was computed as the difference between the RT for Go trials that followed incorrect Stop trials and Go trials that immediately preceded incorrect Stop trials on a pairwise basis. The mean of these differences was computed across trials for each participant.^47^

### BOLD fMRI data analysis

On the first-level, for each participant, preprocessed functional images were entered into a voxel-wise general linear model (GLM). Task events were modeled as regressors of interest based on the onsets of the go-signals and convolved with the canonical hemodynamic response function (HRF). Separate regressors were specified for successful go trials (SGo), failed go trials (FGo), successful stop trials (SS), and failed stop trials (FS). Six realignment parameters derived during preprocessing were included as nuisance regressors. A high-pass filter (128 s) was applied to remove low-frequency signal drifts, and temporal autocorrelations were modeled using an autoregressive AR(1) process implemented in SPM12. For each subject, we computed contrast images representing neural activity during successful response inhibition (Successful Stop [SS] > Successful Go [SGo]), failed response inhibition (Failed Stop [FS] > Successful Go [SGo]), and neural activity specifically associated with successful relative to failed stopping (Successful Stop [SS] > Failed Stop [FS]).

On the second level, first, one-sample t-tests were conducted for each contrast to verify the expected task-related activation patterns.^34,42,52^ Second, we examined associations between DSM-5 AUD symptom domains and general neural activation during successful response inhibition (SS > SGo), and activation specifically associated with successful relative to failed stopping (SS > FS), using whole-brain multiple regression analyses in SPM12. Symptom counts for the impaired control, risky use, social problems, and physical dependence domains were entered simultaneously as predictors, while controlling for site-related variability and head motion. To assess whether observed associations reflected overall AUD severity rather than specific symptom domains, additional whole-brain regression analyses were conducted using total DSM-5 AUD symptom count as predictor. Furthermore, a full-factorial model including group (AUD vs. controls) as a between-subject factor was performed to evaluate whether neural activation differences were explained by diagnostic status. Statistical significance was determined using a voxel-wise cluster-defining threshold of *p* < .001 (uncorrected), corrected to a cluster-extent threshold of *p_FWE_*< 0.05.

### Statistical analysis

Associations between DSM-5 AUD symptom domains and behavioral measures of SST performance as well as questionnaire-based drinking outcomes were examined using multiple linear regression models. For EMA-derived drinking outcomes, logistic and linear mixed-effects models with participant-specific random intercepts were computed for categorial (drinking frequency, binge-drinking frequency) and continuous outcomes (amount of drinking), respectively. Symptom counts for the impaired control, risky use, social problems, and physical dependence domains were entered simultaneously to estimate their unique associations while controlling for the remaining symptom domains. Models were adjusted for age, sex, and study site. Variance inflation factor (VIF) analysis indicated no evidence of problematic multicollinearity (**Tab. S1**). To account for multiple testing, *p*-values were FDR-corrected for the number of measures within each outcome domain, including SST-derived behavioral measures (*n* = 3), retrospective self-reported drinking measures (*n* = 3), and EMA-derived drinking measures (*n* = 3).

## Results

### Sample description

Following quality control procedures, the final sample comprised 589 participants spanning a broad range of AUD symptom severity, including 432 individuals meeting DSM-5 criteria for AUD and 157 controls. Demographic, clinical, and task-related characteristics are summarized in **Tab. 1**.

**Table 1.**
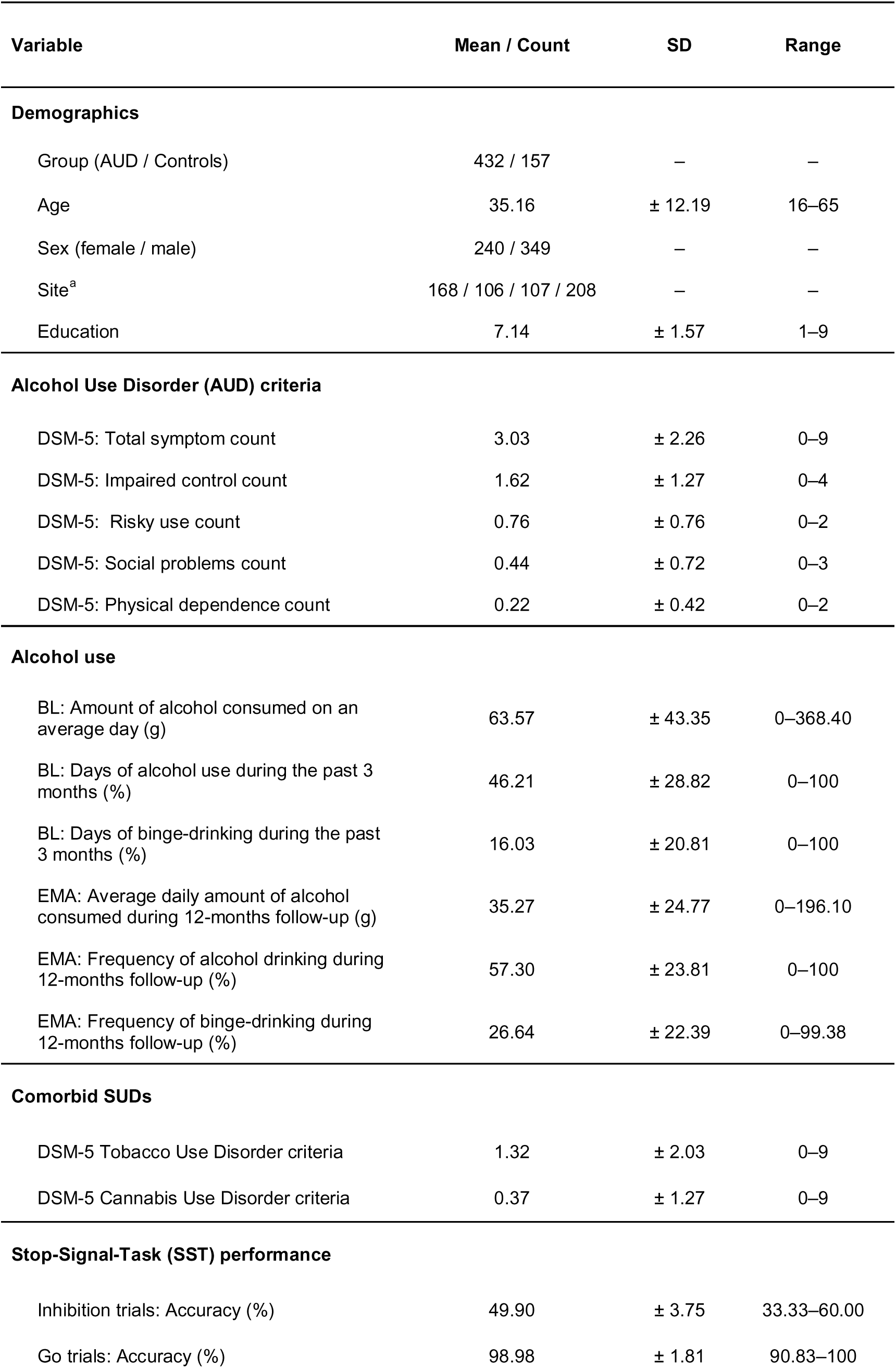
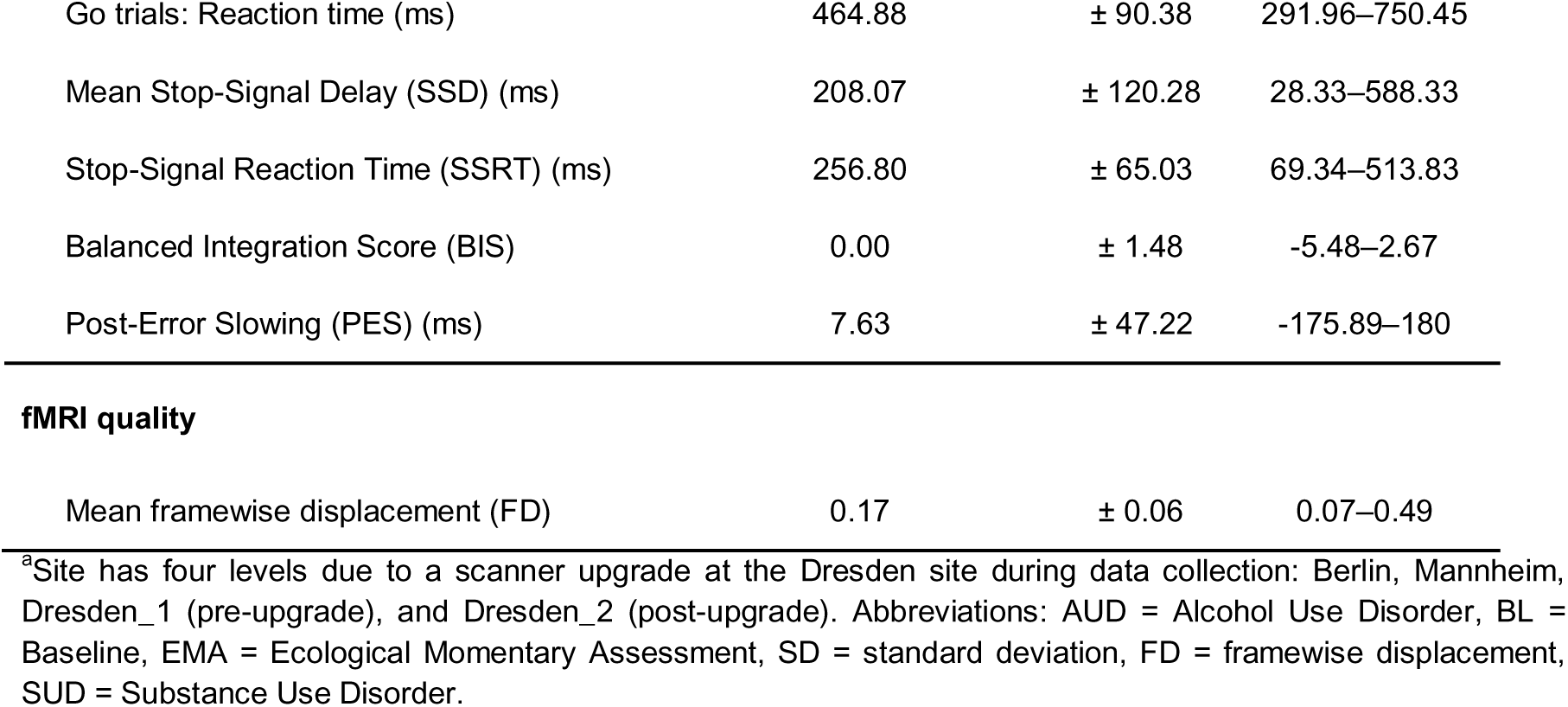
Sample description and demographics (*N* = 589).

| Variable | Mean / Count | SD | Range |
| --- | --- | --- | --- |
| **Demographics** |  |  |  |
| Group (AUD / Controls) | 432 / 157 | – | – |
| Age | 35.16 | $\pm 12.19$ | 16–65 |
| Sex (female / male) | 240 / 349 | – | – |
| Site<sup>a</sup> | 168 / 106 / 107 / 208 | – | – |
| Education | 7.14 | $\pm 1.57$ | 1–9 |
| **Alcohol Use Disorder (AUD) criteria** |  |  |  |
| DSM-5: Total symptom count | 3.03 | $\pm 2.26$ | 0–9 |
| DSM-5: Impaired control count | 1.62 | $\pm 1.27$ | 0–4 |
| DSM-5: Risky use count | 0.76 | $\pm 0.76$ | 0–2 |
| DSM-5: Social problems count | 0.44 | $\pm 0.72$ | 0–3 |
| DSM-5: Physical dependence count | 0.22 | $\pm 0.42$ | 0–2 |
| **Alcohol use** |  |  |  |
| BL: Amount of alcohol consumed on an average day (g) | 63.57 | $\pm 43.35$ | 0–368.40 |
| BL: Days of alcohol use during the past 3 months (%) | 46.21 | $\pm 28.82$ | 0–100 |
| BL: Days of binge-drinking during the past 3 months (%) | 16.03 | $\pm 20.81$ | 0–100 |
| EMA: Average daily amount of alcohol consumed during 12-months follow-up (g) | 35.27 | $\pm 24.77$ | 0–196.10 |
| EMA: Frequency of alcohol drinking during 12-months follow-up (%) | 57.30 | $\pm 23.81$ | 0–100 |
| EMA: Frequency of binge-drinking during 12-months follow-up (%) | 26.64 | $\pm 22.39$ | 0–99.38 |
| **Comorbid SUDs** |  |  |  |
| DSM-5 Tobacco Use Disorder criteria | 1.32 | $\pm 2.03$ | 0–9 |
| DSM-5 Cannabis Use Disorder criteria | 0.37 | $\pm 1.27$ | 0–9 |
| **Stop-Signal-Task (SST) performance** |  |  |  |
| Inhibition trials: Accuracy (%) | 49.90 | $\pm 3.75$ | 33.33–60.00 |
| Go trials: Accuracy (%) | 98.98 | $\pm 1.81$ | 90.83–100 |
| Go trials: Reaction time (ms) | 464.88 | ± 90.38 | 291.96–750.45 |
| Mean Stop-Signal Delay (SSD) (ms) | 208.07 | ± 120.28 | 28.33–588.33 |
| Stop-Signal Reaction Time (SSRT) (ms) | 256.80 | ± 65.03 | 69.34–513.83 |
| Balanced Integration Score (BIS) | 0.00 | ± 1.48 | -5.48–2.67 |
| Post-Error Slowing (PES) (ms) | 7.63 | ± 47.22 | -175.89–180 |
| **fMRI quality** |  |  |  |
| Mean framewise displacement (FD) | 0.17 | ± 0.06 | 0.07–0.49 |

<sup>a</sup>Site has four levels due to a scanner upgrade at the Dresden site during data collection: Berlin, Mannheim, Dresden\_1 (pre-upgrade), and Dresden\_2 (post-upgrade). Abbreviations: AUD = Alcohol Use Disorder, BL = Baseline, EMA = Ecological Momentary Assessment, SD = standard deviation, FD = framewise displacement, SUD = Substance Use Disorder.

### Behavioral inhibition performance by AUD symptom domain

After covariate control, a higher symptom count in the impaired control domain was associated with poorer overall task performance on go trials, as indicated by lower Balanced Integration Scores (BIS; β = −0.12, 95% CI = [-0.22; −0.03], *p*_FDR_ = .026, *f*² = 0.01), and reduced behavioral adaptation following errors, reflected in diminished post-error slowing (PES; β = −0.12, 95% CI = [-0.23; −0.02], *p*_FDR_ = .026, *f*² = 0.01). No significant association was observed with stop-signal reaction times (SSRTs) after adjustment for covariates. Moreover, no significant associations were found between any of the other AUD symptom domains (risky use, social problems, or physical dependence) and behavioral performance measures during the SST (**Fig. 1, Tab. S2**).

**Figure 1.**
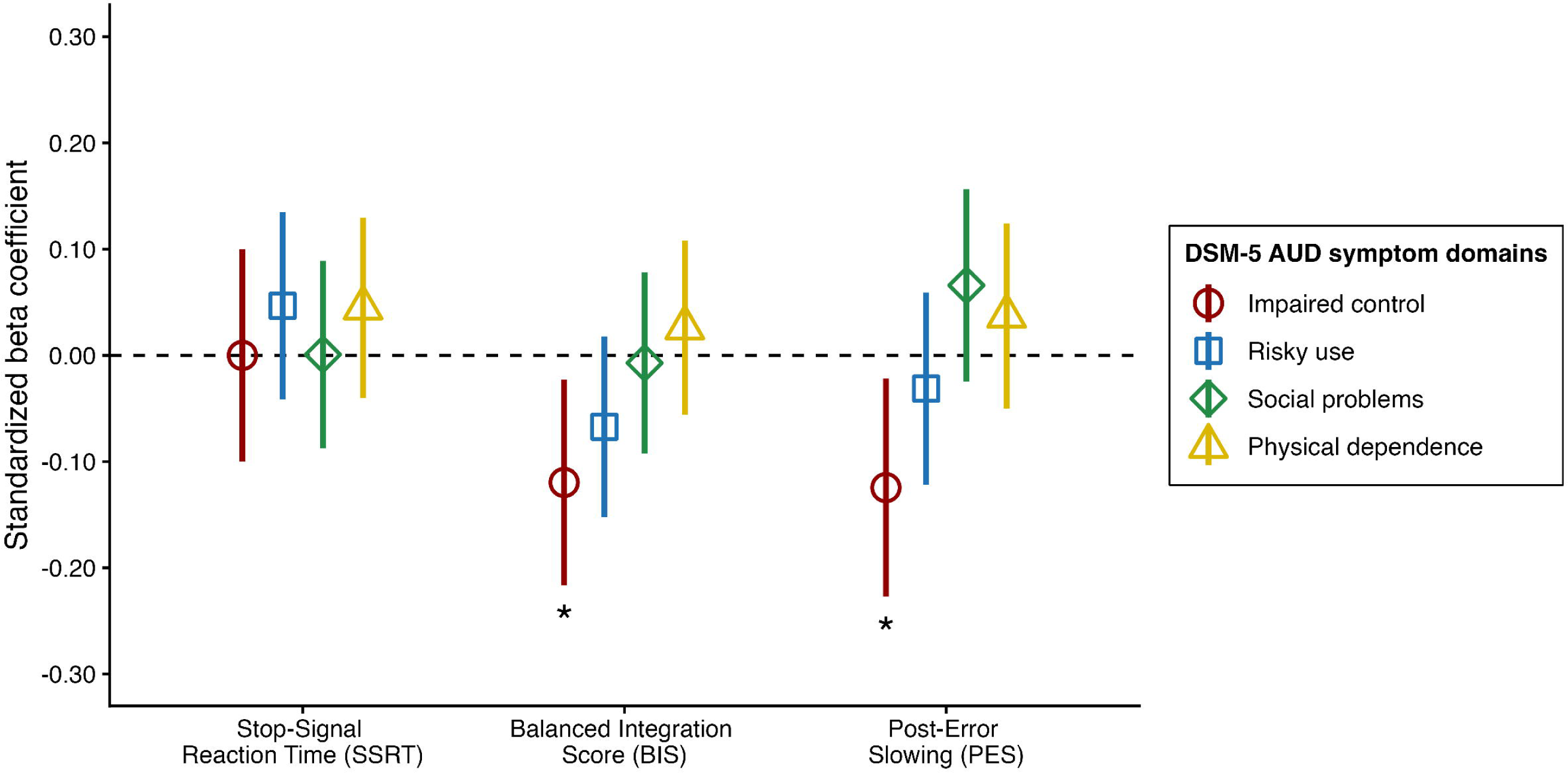
Associations between Stop-Signal Task (SST) performance measures and DSM-5 Alcohol Use Disorder (AUD) symptom domains. Greater symptom severity in the impaired control domain was associated with reduced speed–accuracy integration, reflected by lower Balanced Integration Scores (BIS), and diminished behavioral adaptation following errors, reflected by reduced post-error slowing (PES). All associations were adjusted for age, sex, and site. \**p* < .05, \*\**p* < .01, \*\*\**p* < .001 (FDR-corrected).

### Neural activation during inhibitory control demands by AUD symptom domain

Whole-brain one-sample t-tests demonstrated the expected task-related activation patterns across all SST contrasts, with engagement of key inhibition-related brain regions (**Fig. S2**). Next, to assess associations of neural activation during inhibitory control with AUD symptom domains, we conducted separate whole-brain multiple regression analyses testing the association between each symptom domain and neural activation during response inhibition, while controlling for the remaining symptom domains, study site, and mean head motion.

For the Successful Stop (SS) > Successful Go (SGo) contrast, reflecting neural activation during response inhibition, greater symptom severity in the impaired control domain was associated with reduced activation in five clusters encompassing (i) the bilateral cerebellum extending into occipital and precuneus regions, (ii) a left-lateralized fronto-insular-parietal cluster including the middle and inferior frontal gyri, insula, and supramarginal gyrus, (iii) a right-lateralized fronto-temporo-insular cluster extending into inferior parietal regions, (iv) the supplementary motor area and dorsal anterior cingulate cortex, and (v) the mid-cingulate cortex (**Fig. 2**). Peak coordinates and corresponding AAL3 labels for each cluster are reported in **Tab. 2**. No positive associations with impaired control symptoms were observed. Higher symptom count in the social impairment domain was associated with increased activation in the parieto-occipital gyrus (**Tab. S3, Fig. 2**). Apart from that, no other significant positive or negative associations were observed for the risky use, social impairment, or physical dependence domains. Importantly, total AUD severity and AUD diagnostic status were not significantly associated with brain activation during inhibition.

**Figure 2.**
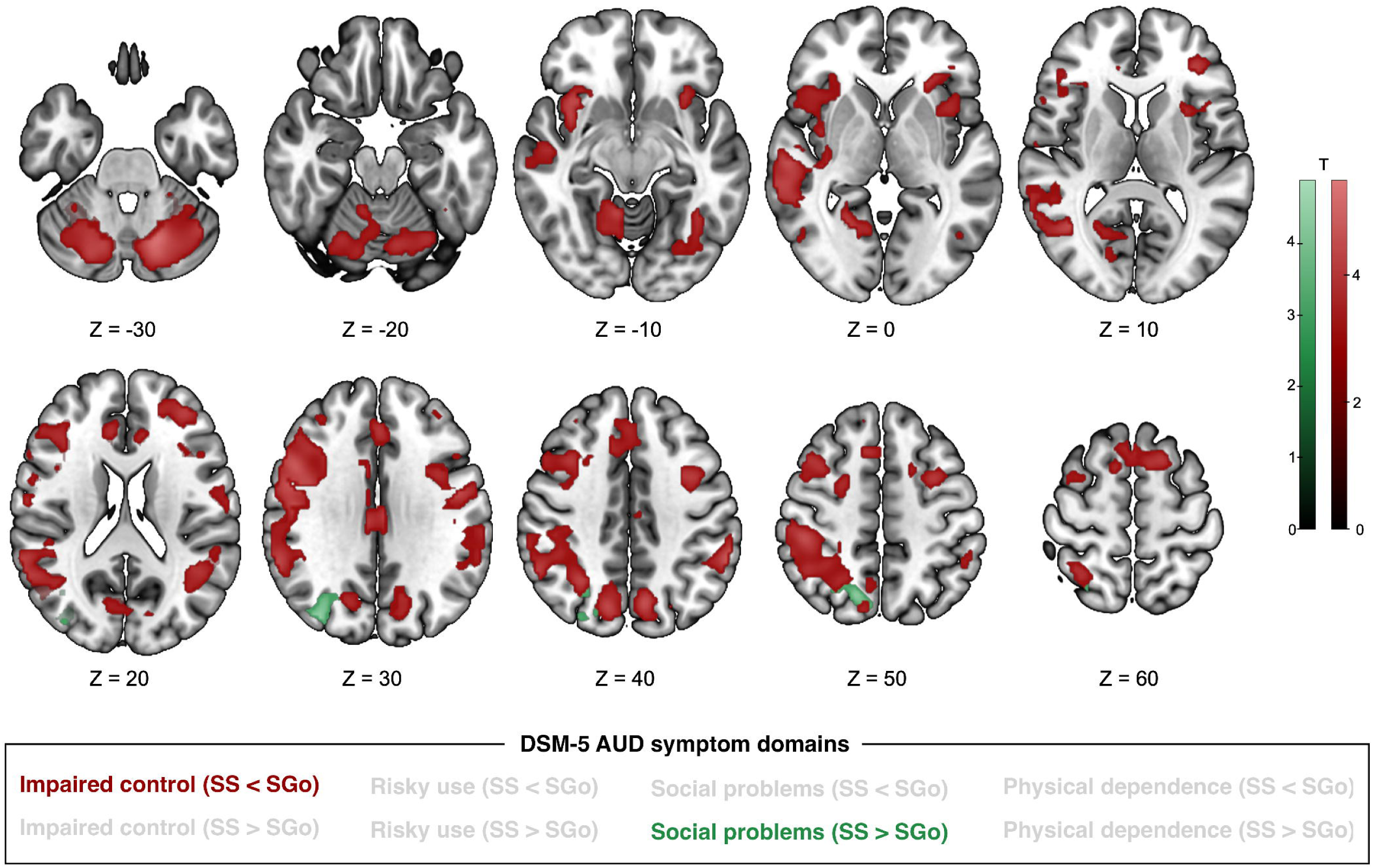
Whole-brain multiple regression analyses examining associations between DSM-5 AUD symptom domains and neural activation during response inhibition (Successful Stop [SS] vs. Successful Go [SGo]). Greater symptom severity in the impaired control domain was associated with widespread reductions in neural activation during response inhibition, whereas no significant associations were observed for risky use or physical dependence symptoms. A small positive association was observed for the social problems domain in a parieto-occipital cluster. Significant activation was determined by a cluster-defining threshold of *p* < 0.001 (uncorrected), corrected to a cluster-extent threshold of *p_FWE_* < 0.05.

**Table 2.**
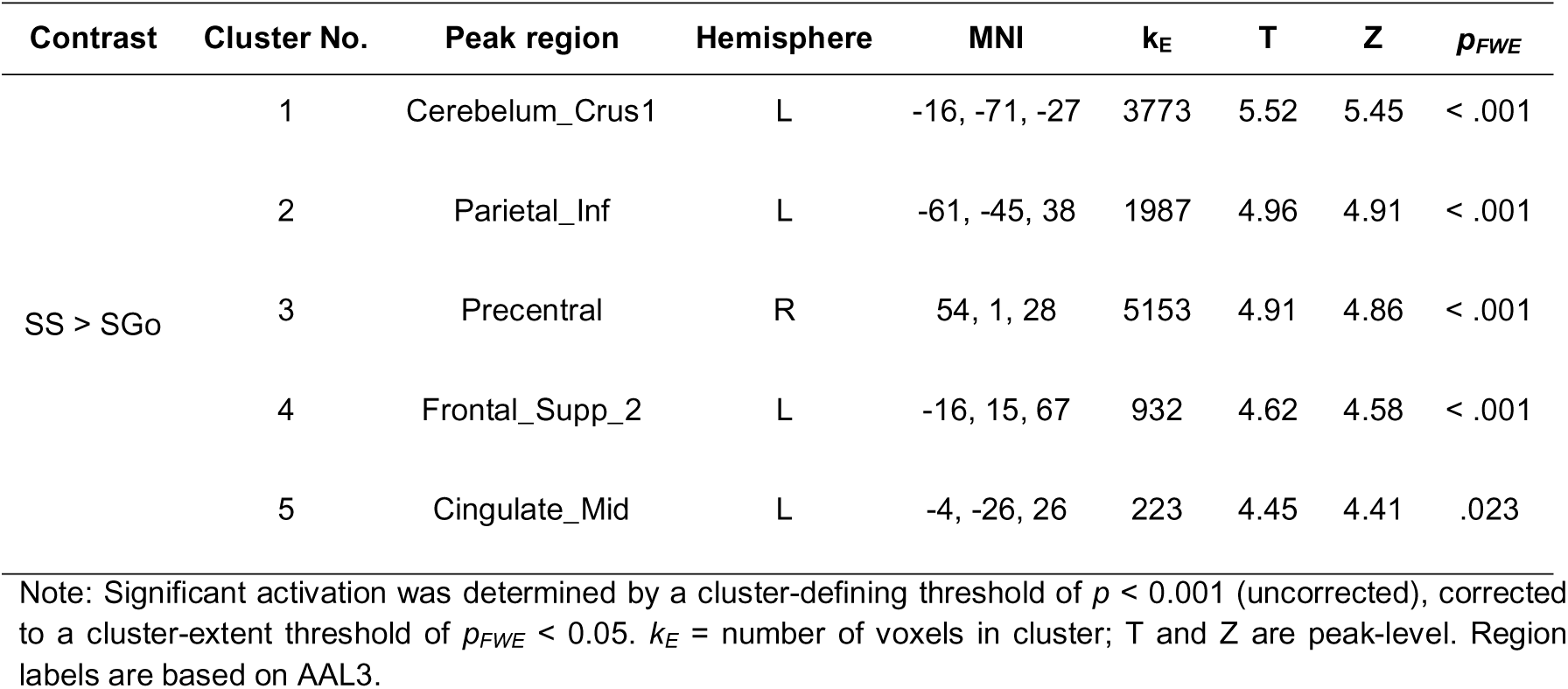
Whole-brain associations between impaired control symptom severity and neural activation during successful response inhibition (SS > SGo), controlling for other AUD symptom domains, site, and mean FD.

For the Successful Stop (SS) > Failed Stop (FS) contrast, reflecting neural processes specifically related to successful inhibitory control, greater symptom severity in the impaired control domain was associated with reduced activation in the right middle/inferior frontal gyrus (peak MNI [42, 25, 24]; *k_E_* = 540; *T* = 4.20; *Z* = 4.17; cluster-level *p_FWE_* = .001; **Fig. 3**). Notably, no significant positive or negative associations were observed for the risky use, social problems, or physical dependence domains. Likewise, neither total AUD symptom count nor AUD diagnostic status was associated with significant activation differences in either direction, suggesting that activation differences during successful inhibitory control are selectively associated with impaired control symptoms.

**Figure 3.**
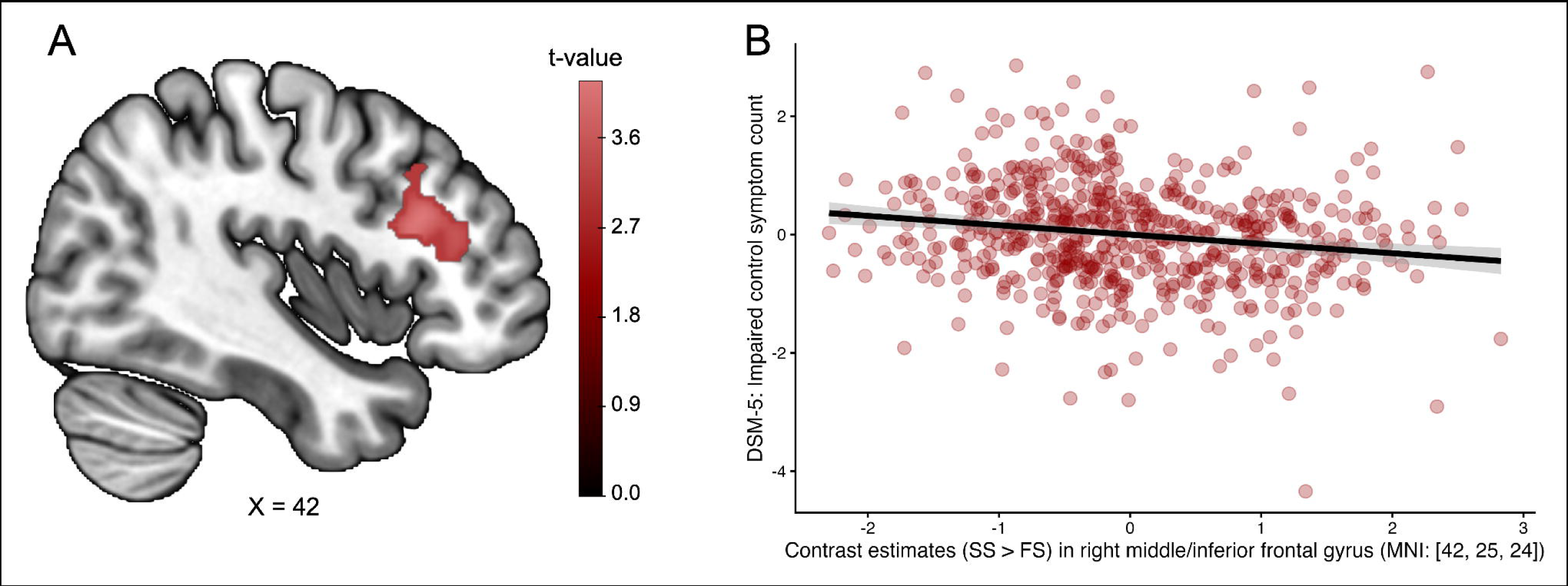
**A)** Whole-brain analysis of the Successful Stop (SS) > Failed Stop (FS) contrast showed that reduced right middle/inferior frontal gyrus (MFG/IFG; peak MNI coordinates: [42, 25, 24]) recruitment during successful inhibitory control was related to greater DSM-5 impaired control symptom severity. Significant activation was determined by a cluster-defining threshold of *p* < 0.001 (uncorrected), corrected to a cluster-extent threshold of *p_FWE_* < 0.05. **B)** Partial regression plot illustrating the association between impaired control symptom count and extracted contrast estimates from the significant right MFG/IFG cluster, after adjustment for other DSM-5 symptom domains (risky use, social problems, physical dependence), study site, and mean framewise displacement.

### Associations with drinking behaviors by AUD symptom domain

For the questionnaire-based drinking measures at baseline, greater severity in the impaired control and risky use symptom domains was associated with higher average daily alcohol consumption, more frequent alcohol use, and more frequent binge drinking, respectively (**Fig. 4, Tab. S4**). For EMA-derived real-life drinking behavior assessed over 12 months in participants with AUD, greater severity in the impaired control domain as well as the physical dependence domain was associated with higher daily alcohol consumption and more frequent binge-drinking episodes (**Fig. 5, Tab. S5**). Greater physical dependence severity was additionally associated with higher drinking frequency. Notably, impaired control was the only symptom domain that demonstrated robust associations with alcohol use across both questionnaire-based and real-life assessments of drinking behavior.

**Figure 4.**
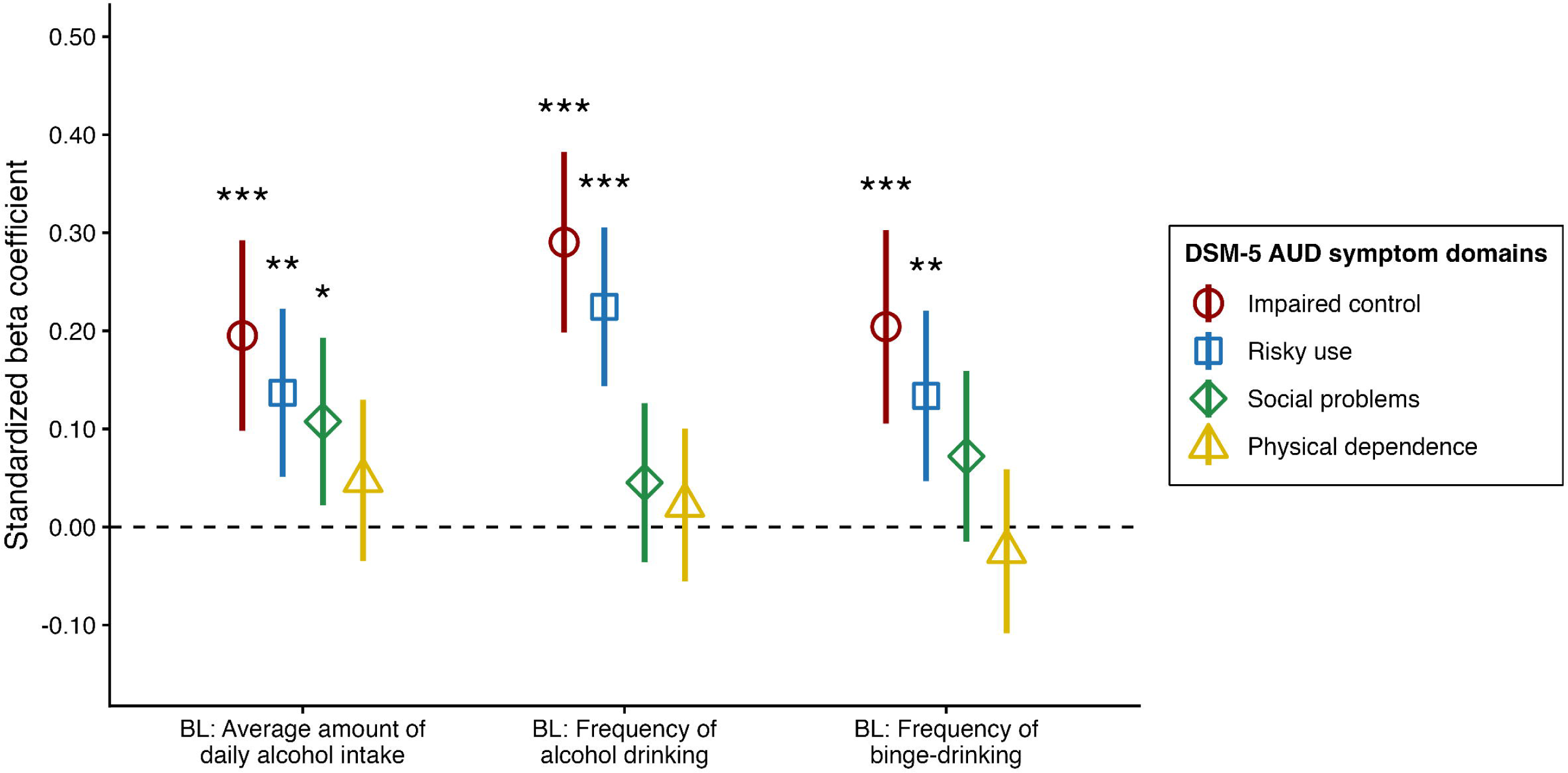
Associations between alcohol drinking behaviors assessed at Baseline (BL) and DSM-5 Alcohol Use Disorder (AUD) symptom domains. Greater symptom severity, particularly in the impaired control and risky use domains, was associated with more severe drinking behaviors, indicated by higher amounts of alcohol consumption, more frequent drinking, and more frequent binge drinking. All associations were adjusted for age, sex, and study site. \**p* < .05, \*\**p* < .01, \*\*\**p* < .001 (FDR-corrected).

**Figure 5.**
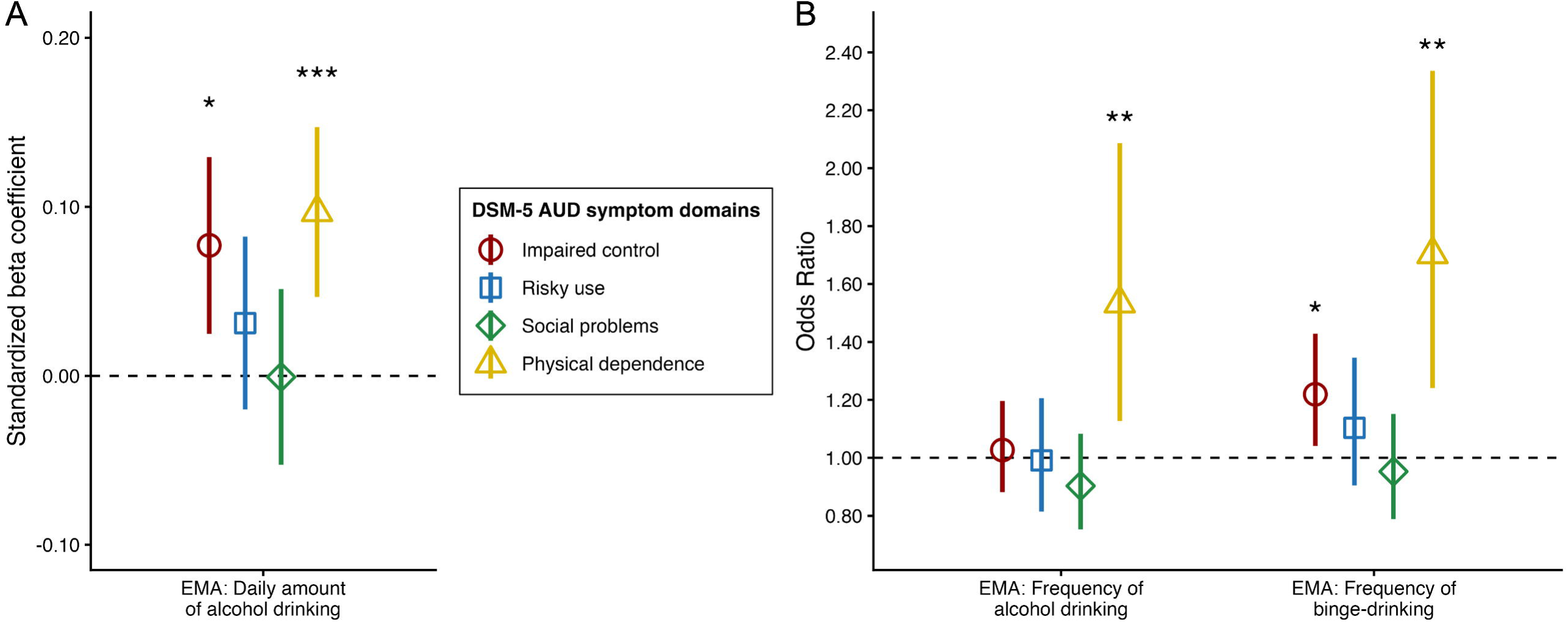
Associations between alcohol drinking behaviors assessed during Ecological Momentary Assessment (EMA) and DSM-5 Alcohol Use Disorder (AUD) symptom domains. A) Greater severity of impaired control and physical dependence symptoms was associated with increased daily alcohol consumption in real-life settings. B) Greater severity of impaired control and physical dependence symptoms was associated with increased odds of binge drinking, while physical dependence symptoms were additionally associated with higher drinking frequency. All associations were adjusted for age, sex, and study site. \**p* < .05, \*\**p* < .01, \*\*\**p* < .001 (FDR-corrected).

### Exploration of associations between neural activation during inhibitory control and drinking behavior

Finally, we explored whether neural activation in regions associated with impaired control symptoms was related to questionnaire-based and EMA-derived measures of drinking behavior. Mean activation estimates were extracted from each cluster and correlated with drinking outcomes. However, none of these associations reached statistical significance (**Tab. S6**).

## Discussion

The present study investigated whether behavioral and neural markers of inhibitory control are differentially associated with distinct DSM-5 symptom domains of alcohol use disorder (AUD). Across a large sample spanning a broad spectrum of AUD symptom severity, we found converging evidence that alterations in inhibitory control were selectively associated with DSM-5 AUD symptoms relating to impaired control, whereas symptom domains covering risky patterns of alcohol use, social impairments, and physical dependence symptoms showed little to no relationship with inhibition-related measures. Specifically, greater impaired control symptom severity was associated with poorer behavioral performance in go trials and diminished post-error adaptation as well as reduced activation in distributed brain regions related to inhibitory control during the SST. Furthermore, impaired control symptomatology was consistently associated with higher alcohol consumption across questionnaire-based and real-world drinking data. Together, our findings suggest that inhibitory control alterations are selectively linked to the impaired control symptom dimension of AUD, rather than AUD in general.

Within the DSM-5 framework, impaired control symptoms reflect failures to regulate alcohol use despite intentions to limit consumption, including unsuccessful attempts to cut down drinking and using alcohol in larger amounts or over longer periods than intended.^31^ Such symptoms therefore possess a conceptual proximity to inhibitory control mechanisms that is largely absent for symptom domains reflecting social consequences, risky use, or physiological dependence. This specificity may help explain inconsistent evidence for inhibitory control deficits in AUD in behavioral and neuroimaging studies.^17–23,29,30^ Previous work has predominantly relied on categorical case-control designs, implicitly assuming relative homogeneity among individuals meeting diagnostic criteria for AUD. However, AUD can emerge from numerous symptom combinations that may reflect partially distinct neurocognitive subtypes.^32–35^ The present findings suggest that alterations in inhibitory control are particularly characteristic of individuals with impaired control symptoms.

At the behavioral level, individuals reporting greater impaired control symptom severity exhibited lower Balanced Integration Scores (BIS) and reduced post-error slowing during the Stop-Signal Task. The BIS integrates information on response speed and accuracy and captures overall task performance.^44,45^ As the BIS is derived from performance on Go trials, this finding may reflect impairments in maintaining goal-directed task performance across repeated actions. Post-error slowing refers to the tendency of prolonged response times after committing an error, and is widely considered a behavioral marker of performance monitoring and adaptive cognitive control.^47,48^ Reduced post-error slowing has previously been reported in individuals with alcohol dependence^14^ and has been found to moderate the relationship between craving and subsequent alcohol use.^49^ In contrast, no significant association was observed for Stop-Signal Reaction Time (SSRT), the most widely used behavioral measure of response inhibition, after adjustment for demographic covariates. Together, these findings suggest that impaired control symptoms are more closely related to reduced efficiency in maintaining goal-directed behavior and adapting performance following errors than to behavioral response inhibition deficits per se. Successful regulation of alcohol use requires individuals to monitor the consequences of their behavior and adjust their actions in accordance with longer-term goals. Difficulties in maintaining effective goal-directed performance and adapting behavior following performance failures may therefore contribute to the persistence of alcohol use despite repeated negative consequences, a defining feature of AUD.

At the neural level, greater impaired control symptom severity was associated with reduced recruitment of fronto-insular, parietal, cingulate, supplementary motor, and cerebellar regions during successful response inhibition, as well as diminished activation of the right inferior/middle frontal gyrus during successful relative to unsuccessful stopping. These regions constitute key nodes of the inhibitory control network and have been consistently implicated in the implementation of response inhibition,^28,50,52^ with the right inferior frontal gyrus, extending into the anterior insula, playing a particularly prominent role in the implementation of stopping processes.^42,51,52^ Notably, these neural alterations emerged selectively in relation to impaired control symptoms and were absent for other symptom dimensions, overall AUD severity, and conventional AUD vs. control group comparisons. This pattern suggests that inhibitory control network dysfunction may not represent a general feature of AUD, but rather a neurocognitive mechanism specifically linked to the loss of control over alcohol use. Consistent with this interpretation, impaired control was also the only symptom domain that showed robust and convergent associations with alcohol consumption patterns. Based on questionnaire data, individuals reporting greater impaired control symptom severity consumed larger amounts of alcohol, drank more frequently, and engaged in binge drinking more often, with comparable associations observed in EMA-based assessments of real-world drinking over a 12-months follow-up period.

The convergence of our findings across neural, behavioral, and ecological data provides evidence that inhibitory control deficits are selectively linked to impaired control symptoms rather than representing a ubiquitous feature of AUD. This distinction may have important implications for treatment development. Despite considerable interest in interventions targeting inhibitory control, such as response inhibition trainings, evidence for their effectiveness in reducing alcohol use has remained mixed, with studies reporting variable effects across outcomes and populations.^53–59^ The current findings raise the possibility that these limited effects may partly reflect a mismatch between intervention targets and patient characteristics. If inhibitory control deficits are primarily associated with impaired control symptoms, interventions designed to strengthen inhibitory control may be most beneficial for individuals exhibiting this specific symptom profile, while providing limited benefit for patients for whom other symptom dimensions are dominant. Therefore, our results may provide novel insights into the potential value of symptom-based approaches in advancing addiction treatment.

Several limitations should be acknowledged when interpreting the results. First, symptom domains were derived from theoretically defined symptom clusters based on the DSM-5.^31^ Alternative dimensional frameworks, such as data-driven clustering of AUD symptoms,^35^ may capture different aspects of clinical heterogeneity and could be examined in future studies. Second, the impaired control symptom cluster exhibits substantially greater between-subject variability than the other symptom domains. This restricted variability may have reduced sensitivity to detect associations with these symptom domains and should therefore be considered when interpreting the apparent specificity of our findings to impaired control symptoms. Third, EMA data were available only for a subset of participants and exclusively among individuals with AUD. Consequently, the extent to which EMA-based findings generalize across the broad continuum of alcohol use severity, including social-drinking controls represented in the other analyses, remains elusive. Fourth, although impaired control symptoms were robustly associated with both questionnaire-based and real-world drinking behavior, the present study did not examine their prospective relationship with other clinically relevant outcomes, such as treatment response, or relapse risk.^60,61^

In sum, our study shows that inconsistent evidence for inhibitory control deficits in AUD may partly reflect the substantial clinical heterogeneity of the disorder, with neural and behavioral alterations becoming detectable primarily when focusing on the DSM-5 symptom dimension that is conceptually linked to inhibitory control itself (i.e., impaired control), while no such aberrations are observed for other symptom domains (social problems, risky use, physical dependence). These findings highlight the potential of symptom-based approaches for understanding heterogeneity in addiction.

## Acknowledgements

This research was supported by the German Research Foundation (Deutsche Forschungsgemeinschaft, DFG, project numbers 402170461 [TRR 265 to HW, MM, KR, MR, FK, PB, BL, AH, TE, MNS, RS], 454245598 [IRTG 2773 to MM, MNS], and 521379614 [TRR 393 to MNS]). We thank the non-author collaborators of the ReCoDe-Consortium (https://www.trr265.org/en). Generative AI tools were used only for minor language editing and to improve readability. All research ideas, study design, experiments, analyses, and interpretations were conceived and carried out by the authors. The authors take full responsibility for the originality, validity, and integrity of the work.

## References

1. World Health Organization (WHO). Global status report on alcohol and health and treatment of substance use disorders 2024. Geneva: World Health Organization; 2024.

2. GBD 2016 Alcohol Collaborators. Alcohol use and burden for 195 countries and territories, 1990-2016: a systematic analysis for the Global Burden of Disease Study 2016. Lancet. 2018;392(10152):1015–1035.

3. Diamond A. Executive functions. Annu Rev Psychol. 2013:64:135–68.

4. Chambers CD, Garavan H, Bellgrove MA. Insights into the neural basis of response inhibition from cognitive and clinical neuroscience. Neurosci Biobehav Rev. 2009;33(5):631–46.

5. Nigg JT. On inhibition/disinhibition in developmental psychopathology: views from cognitive and personality psychology and a working inhibition taxonomy. Psychol Bull. 2000;126(2):220–46.

6. López-Caneda E, Rodrígez Holguín S, Cadaveira F, Corral M, Doallo S. Impact of alcohol use on inhibitory control (and vice versa) during adolescence and young adulthood: a review. Alcohol Alcohol. 2014;49(2):173–81.

7. Norman AL, Pulido C, Squeglia LM, Spadoni AD, Paulus MP, Tapert SF. Neural Activation during Inhibition Predicts Initiation of Substance Use in Adolescence. Drug Alcohol Depend. 2011;119(3):216–223.

8. Bernard L, Cyr L, Bonnet-Suard A, Cutarella C, Bréjard V. Drawing alcohol craving process: A systematic review of its association with thought suppression, inhibition and impulsivity. Heliyon. 2021;7(1):e05868.

9. Field M, Wiers RW, Christiansen P, Fillmore MT, Verster JC. Acute Alcohol Effects on Inhibitory Control and Implicit Cognition: Implications for Loss of Control Over Drinking. Alcohol Clin Exp Res. 2010;34(8):1346–1352.

10. Moeller SJ, Bederson L, Alia-Klein N, Goldstein RZ. Neuroscience of inhibition for addiction medicine: From prediction of initiation to prediction of relapse. Prog Brain Res. 2015;223:165–188.

11. Littman R, Takács A, Do all inhibitions act alike? A study of go/no-go and stop-signal paradigms. PLoS One. 2017;12(10):e0186774.

12. Raud L, Westerhausen R, Dooley N, Huster RJ. Differences in unity: The go/no-go and stop signal tasks rely on different mechanisms. Neuroimage. 2020;210:116582.

13. Hu S, Ide JS, Zhang S, Sinha R, Li CR. Conflict anticipation in alcohol dependence — A model-based fMRI study of stop signal task. NeuroImage: Clin. 2015;8:39–50.

14. Lawrence AJ, Luty J, Bogdan NA, Sahakian BJ, Clark L. Impulsivity and response inhibition in alcohol dependence and problem gambling. Psychopharmacology. 2009;207(1):163–72.

15. Smith JL, Mattick RP, Jamadar SD, Iredale JM. Deficits in behavioural inhibition in substance abuse and addiction: A meta-analysis. Drug Alcohol Depend. 2014;145:1–33.

16. Wang D, Tian Y, Yang C, Luo Y, Zhao B, Wang G, et al. Post-error slowing predicts for relapse in individuals with alcohol use disorder. J Psychiatr Res. 2023;160:205–212.

17. Bednarski SR, Erdman E, Luo X, Zhang S, Hu S, Li SCR. Neural Processes of an Indirect Analog of Risk Taking in Young Nondependent Adult Alcohol Drinkers—An fMRI Study of the Stop Signal Task. Alcohol Clin Exp Res. 2012;36(5):768–779.

18. Czapla M, Baeuchl C, Simon JJ, Richter B, Kluge M, Friederich HJ et al. Do alcohol-dependent patients show different neural activation during response inhibition than healthy controls in an alcohol-related fMRI go/no-go-task? Psychopharmacology. 2017;234:1001–1015.

19. Fascher M, Nowaczynski S, Spindler C, Strobach T, Muehlhan M. Neural underpinnings of response inhibition in substance use disorders: weak meta-analytic evidence for a widely used construct. Psychopharmacology. 2024;241(1):1–17.

20. Gerhardt S, Luderer M, Bumb JM, Sobanski E, Moggi F, Kiefer F, Vollstädt-Klein S. Stop What You’re Doing!-An fMRI Study on Comparisons of Neural Subprocesses of Response Inhibition in ADHD and Alcohol Use Disorder. Front Psychiatry. 2021;12:691930.

21. Li CR, Luo X, Yan P, Bergquist K, Sinha R. Altered Impulse Control in Alcohol Dependence: Neural Measures of Stop Signal Performance. Alcohol Clin Exp Res. 2009;33(4):740–750.

22. Sjoerds Z, van den Brink W, Beekman ATF, Penninx BWJH, Veltman DJ. Response inhibition in alcohol-dependent patients and patients with depression/anxiety: a functional magnetic resonance imaging study. Psychol Med. 2014;44(8):1713–25.

23. Luijten M, Machielsen MWJ, Veltman DJ, Hester R, de Haan L, Franken IHA. Systematic review of ERP and fMRI studies investigating inhibitory control and error processing in people with substance dependence and behavioural addictions. J Psychiatry Neurosci. 2014;39(3):149–69.

24. Alderson-Myers AB, Arienzo D, Molnar SM, Marinkovic K. Local and network-level dysregulation of error processing is associated with binge drinking. NeuroImage: Clin. 2021;32:102879.

25. Campanella S, Absil J, Carbia Sinde C et al. Neural correlates of correct and failed response inhibition in heavy versus light social drinkers: an fMRI study during a go/no-go task by healthy participants. Brain Imaging Behav. 2017;11:1796–1811.

26. Grieder M, Soravia LM, Tschuemperlin RM et al. Right inferior frontal activation during alcohol-specific inhibition increases with craving and predicts drinking outcome in alcohol use disorder. Front Psych. 2022;13:909992.

27. Nestor LJ, Paterson LM, Murphy A et al. Naltrexone differentially modulates the neural correlates of motor impulse control in abstinent alcohol-dependent and polysubstance-dependent individuals. Eur J Neurosci. 2019;50:2311–2321.

28. Zilverstand A, Huang AS, Alia-Klein N, Goldstein RZ. Neuroimaging Impaired Response Inhibition and Salience Attribution in Human Drug Addiction: A Systematic Review. Neuron. 2018;98(5):886–903.

29. Le TM, Potvin S, Zhornitsky S, Li CR. Distinct patterns of prefrontal cortical disengagement during inhibitory control in addiction: A meta-analysis based on population characteristics. Neurosci Biobehav Rev. 2021;127:255–269.

30. Qiu Z, Wang J. Altered neural activities during response inhibition in adults with addiction: a voxel-wise meta-analysis. Psychol Med. 2021;51(3):387–399.

31. American Psychiatric Association (APA). Diagnostic and statistical manual of mental disorders. 5th ed. Arlington (VA): American Psychiatric Publishing; 2013.

32. Bailey AJ, Ingram PF, Howe LK, Finn PR. Is lower severity alcohol use disorder qualitatively different than more severe manifestations? An evaluation of multivariate symptom clusters. Addiction. 2022;117(6):1598–1608.

33. Brucar LR, Rawls E, Zilverstand A. Neurobiological subtypes in alcohol use disorder and their phenotypic and clinical profiles. Mol Psychiatry. 2026.

34. Hildebrandt MK, Scharz K, Dieterich R, Endrass T. Dissociating the Link of Neural Correlates of Inhibition to the Degree of Substance Use and Substance-Related Problems: A Preregistered, Multimodal, Combined Cross-sectional and Longitudinal Study. Biol Psychiatry. 2023;94(11):898–905.

35. Watts AL, Boness CL, Loeffelman JE, Steinley D, Sher KJ. Does crude measurement contribute to observed unidimensionality of psychological constructs? A demonstration with DSM-5 alcohol use disorder. J Abnorm Psychol. 2021;130(5):512–524.

36. Congdon E, Mumford JA, Cohen JR, Galvan A, Canli T, Poldrack RA. Measurement and reliability of response inhibition. Front Psychol. 2012;3:37.

37. Lipszyc J, Schachar R. Inhibitory control and psychopathology: a meta-analysis of studies using the stop signal task. J Int Neuropsychol Soc. 2010;16(6):1064–76.

38. Verbruggen F, Logan GD. Response inhibition in the stop-signal paradigm. Trends Cogn Sci. 2008;12(11):418–24.

39. Heinz A, Kiefer F, Smolka MN, Endrass T, Beste C, Beck A et al. Addiction Research Consortium: Losing and regaining control over drug intake (ReCoDe)-From trajectories to mechanisms and interventions. Addict Biol. 2020;25(2):e12866.

40. Spanagel R, Bach P, Banaschewski T, Beck A, Bermpohl F, Bernardi RE et al. The ReCoDe addiction research consortium: Losing and regaining control over drug intake-Findings and future perspectives. Addict Biol. 2024;29(7):e13419.

41. Deeken F, Reichert M, Zech H, Wenzel J, Wedemeyer F, Aguilera A et al. Patterns of Alcohol Consumption Among Individuals With Alcohol Use Disorder During the COVID-19 Pandemic and Lockdowns in GermanyJAMA Netw Open. 2022;5(8):e2224641.

42. Gan G, Guevara A, Marxen M, Neumann M, Jünger E, Kobiella A, Mennigen E, Pilhatsch M, Schwarz D, Zimmermann US, Smolka MN. Alcohol-induced impairment of inhibitory control is linked to attenuated brain responses in right fronto-temporal cortex. Biol Psychiatry. 2014;76(9):698–707.

43. Verbruggen F, Aron AR, Band GP, Beste C, Bissett PG, Brockett AT et al. A consensus guide to capturing the ability to inhibit actions and impulsive behaviors in the stop-signal task. Elife. 2019;8:e46323.

44. Liesefeld HR, Fu X, Zimmer HD. Fast and careless or careful and slow? Apparent holistic processing in mental rotation is explained by speed-accuracy trade-offs. J Exp Psychol Learn Mem Cogn. 2015;41(4):1140–1151.

45. Liesefeld HR, Janczyk M. Combining speed and accuracy to control for speed-accuracy trade-offs(?). Behav Res Methods. 2019;51(1):40–60.

46. Rabbitt PM, Errors and error correction in choice-response tasks. J Exp Psychol. 1966;71(2):264–72.

47. Dutilh G, van Ravenzwaaij D, Nieuwenhuis S, van der Maas HLJ, Forstmann BU, Wagenmakers EJ. How to measure post-error slowing: A confound and a simple solution. J Math Psychol. 2012;56(3):208–216.

48. Danielmeier C, Ullsperger M. Post-Error Adjustments. Front Psychol. 2011;2:233.

49. Bresin K, Verona E. Craving and substance use: Examining psychophysiological and behavioral moderators. Int J Psychophysiol. 2021;163:92–103.

50. Isherwood S, Kemp SA, Miletic S, Stevenson N, Bazin PL, Forstmann B. Multi-study fMRI outlooks on subcortical BOLD responses in the stop-signal paradigm. Elife. 2025;12:RP88652.

51. Aron AR, Fletcher PC, Bullmore ET, Sahakian BJ, Robbins TW. Stop-signal inhibition disrupted by damage to right inferior frontal gyrus in humans. Nat Neurosci. 2003;6(2):115–6.

52. Zhang R, Geng X, Lee TMC. Large-scale functional neural network correlates of response inhibition: an fMRI meta-analysis. Brain Struct Funct. 2017;222(9):3973–3990.

53. Houben K, Nederkoorn C, Wiers RW, Jansen A. Resisting temptation: decreasing alcohol-related affect and drinking behavior by training response inhibition. Drug Alcohol Depend. 2011;116:132–136.

54. Bartsch AL, Kothe E, Allom V, Mullan B, Houben K. The effect of non-specific response inhibition training on alcohol consumption: an intervention. J Addict Res Ther. 2016;7:260.

55. Batschelet HM, Stein M, Tschuemperlin RM, Soravia LM, Moggi F. Alcohol-Specific Computerized Interventions to Alter Cognitive Biases: A Systematic Review of Effects on Experimental Tasks, Drinking Behavior, and Neuronal Activation. Front Psychiatry. 2020;10:871.

56. Schenkel EJ, Schöneck R, Wiers RW, Veling H, Becker ES, Lindenmeyer J, Rinck M. Does selective inhibition training reduce relapse rates when added to standard treatment of alcohol use disorder? A randomized controlled trial. Alcohol Clin Exp Res. 2023;47(5):963–974.

57. Reichl D, Enewoldsen N, Müller A, Steins-Loeber S. Pilot testing of an adaptive, individualized inhibitory control training for binge drinking: first evidence on feasibility, acceptance, and efficacy. Psychol Res. 2023;87(4):1267–1283.

58. Stein M, Soravia LM, Tschuemperlin RM, Batschelet HM, Jaeger J, Roesner S, Keller A, Penedo JMG, Wiers RW, Moggi F. Alcohol-specific inhibition training in patients with alcohol use disorder: a multi-centre, double-blind randomized clinical trial examining drinking outcome and working mechanisms. Addiction. 2023;118(4):646–657.

59. Guarriello A, Fleckenstein T, Soravia LM, Tschuemperlin RM, Batschelet HM, Jaeger J, Wiers RW, Moggi F, Stein M. Mediation of beneficial effects of an alcohol-specific inhibition training on drinking of patients with alcohol use disorder: The role of cognitive demands and inhibitory performance. Addict Behav. 2025;161:108212.

60. Böhmer J, Reinhardt P, Garbusow M, Marxen M, Smolka MN, Zimmermann US, Heinz A, Bzdok D, Friedel E, Kruschwitz JD, Walter H. Aberrant functional brain network organization is associated with relapse during 1-year follow-up in alcohol-dependent patients. Addict Biol. 2023;28(11):e13339.

61. Böhmer J, Marxen M, Veer IM, Garbusow M, Bottino M, Zimmermann US, Smolka MN, Heinz A, Friedel E, Heim C, Kruschwitz JD, Walter H. Relapse in alcohol dependence is characterized by disrupted modular brain network organization. Eur Arch Psychiatry Clin Neurosci. 2026;276(4):1733–1748.

