## Supplementary Information for "Impaired Control Symptoms in Alcohol Use Disorder Show Convergent Associations Across Behavioral and Neural Markers of Inhibitory Control and Drinking Behavior"

**Table of contents**

Extended Methods

Figure S1. Diagnostic criteria for Alcohol Use Disorder (AUD) based on DSM-5, and their correspondence to overarching symptom domains.

Figure S2. Task-elicited activation based on one-sample t-tests (*n* = 589) from contrasts A) Successful Stop > Successful Go; B) Failed Stop > Successful Go; and C) Successful Stop > Failed Stop.

Table S1. Variance inflation factors (VIFs) for all predictors included in the linear regression models, calculated to assess potential multicollinearity.

Table S2. Associations between behavioral inhibitory control performance parameters in Stop-Signal Task (SST) and AUD symptom domains in DSM-5 (n = 589).

Table S3. Whole-brain associations between social problems symptom severity and neural activation during successful response inhibition (SS > SGo), controlling for other AUD symptom domains, age, site, and mean FD.

Table S4. Associations between alcohol drinking behaviors assessed at Baseline (BL) and AUD symptom domains in DSM-5 (n = 589).

Table S5. Associations between alcohol drinking behaviors assessed during 12-months Ecological Momentary Assessment (EMA) and AUD symptom domains in DSM-5 (n = 389).

Table S6. Exploration of associations between neural activation in clusters associated with impaired control symptoms during inhibitory control demands and drinking behavior.

**Extended Methods**

Experimental design – The Stop-Signal Task (SST)

During go trials, the go stimulus was presented until the participant responded, capped at a maximum of 1000 ms. During stop trials, the go stimulus remained visible for the length of the adaptive SSD. The subsequent stop signal was then presented for 300 ms on successful stop trials or until a response occurred on failed stop trials. Each trial was followed by a fixation cross displayed for a jittered intertrial interval with a mean duration of 900 ms (range: 700–1100 ms). Go stimuli pointing left or right were presented with equal frequency in both go and stop conditions. Altogether 300 trials were presented, with 20% stop trials. The SST was administered using Presentation® software (Neurobehavioral Systems, Albany, California).

fMRI data acquisition

MRI data were collected on 3T Siemens Prisma/Prisma Fit and Trio scanners in Berlin, Dresden and Mannheim, with a 64-channel coil in Berlin and Mannheim, and a 32-channel coil in Dresden. Functional images were collected with a multi-band T2*-weighted EPI sequence (500 volumes; TR = 869 ms; TE = 38 ms; 60 slices; slice thickness = 2.4 mm; voxel size = 2.4 × 2.4 × 2.4 mm; no slice gap; FoV = 210 mm; matrix = 88 × 88; acquisition orientation T > C; interleaved order; multiband factor = 6; flip angle = 58°; bandwidth = 1832 Hz/Px; prescan normalize; weak raw data filter; fat saturation). Field maps were obtained with a Siemens dual gradient-echo sequence (TR = 698 ms; TE1 = 5.19 ms; TE2 = 7.65 ms; 64 slices; slice thickness = 2.4 mm; no gap; voxel size = 2.4 × 2.4 × 2.4 mm; FoV = 210 mm; matrix = 88 × 88; orientation T > C; interleaved order; flip angle = 54°; bandwidth = 279 Hz/Px). Structural T1-weighted images (MPRAGE) for normalization were acquired with TR = 2000 ms, TE = 2.01 ms, TI = 880 ms, FoV = 256 mm, 208 sagittal slices, 1 × 1 × 1 mm voxels, flip angle = 8°, GRAPPA factor 2 (PE), prescan normalize, 23.1% slice oversampling, and bandwidth = 240 Hz/Px.

fMRI data preprocessing

Preprocessing of functional images was conducted using SPM12, and included slice-timing correction, realignment, and B0-fieldmap correction. T1 images were coregistrated with the EPI images, and normalized to MNI space. EPI images were normalized to MNI152 standard space at 2.4 mm resolution using the transform obtained from the T1 normalization. Spatial smoothing was applied with an 8mm FWHM Gaussian kernel. All images were quality checked by visual inspection. Participants exhibiting insufficient fMRI data quality were excluded from further analyses. Moreover, subjects with excessive head motion (mean framewise displacement (*FD*) > 0.5 mm) were excluded (*n* = 2).

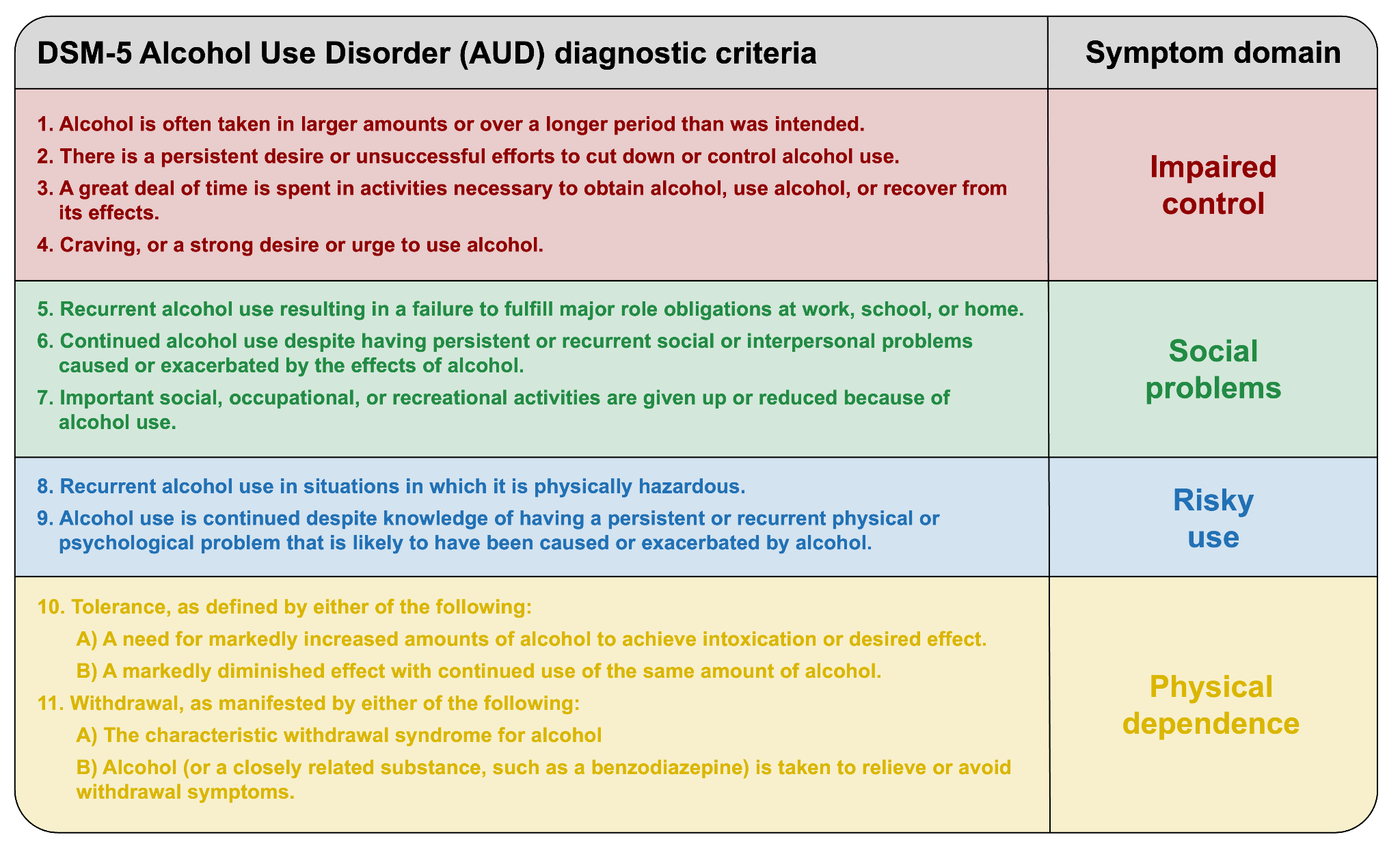

**Figure S1**. Diagnostic criteria for Alcohol Use Disorder (AUD) based on DSM-5, and their correspondence to overarching symptom domains.

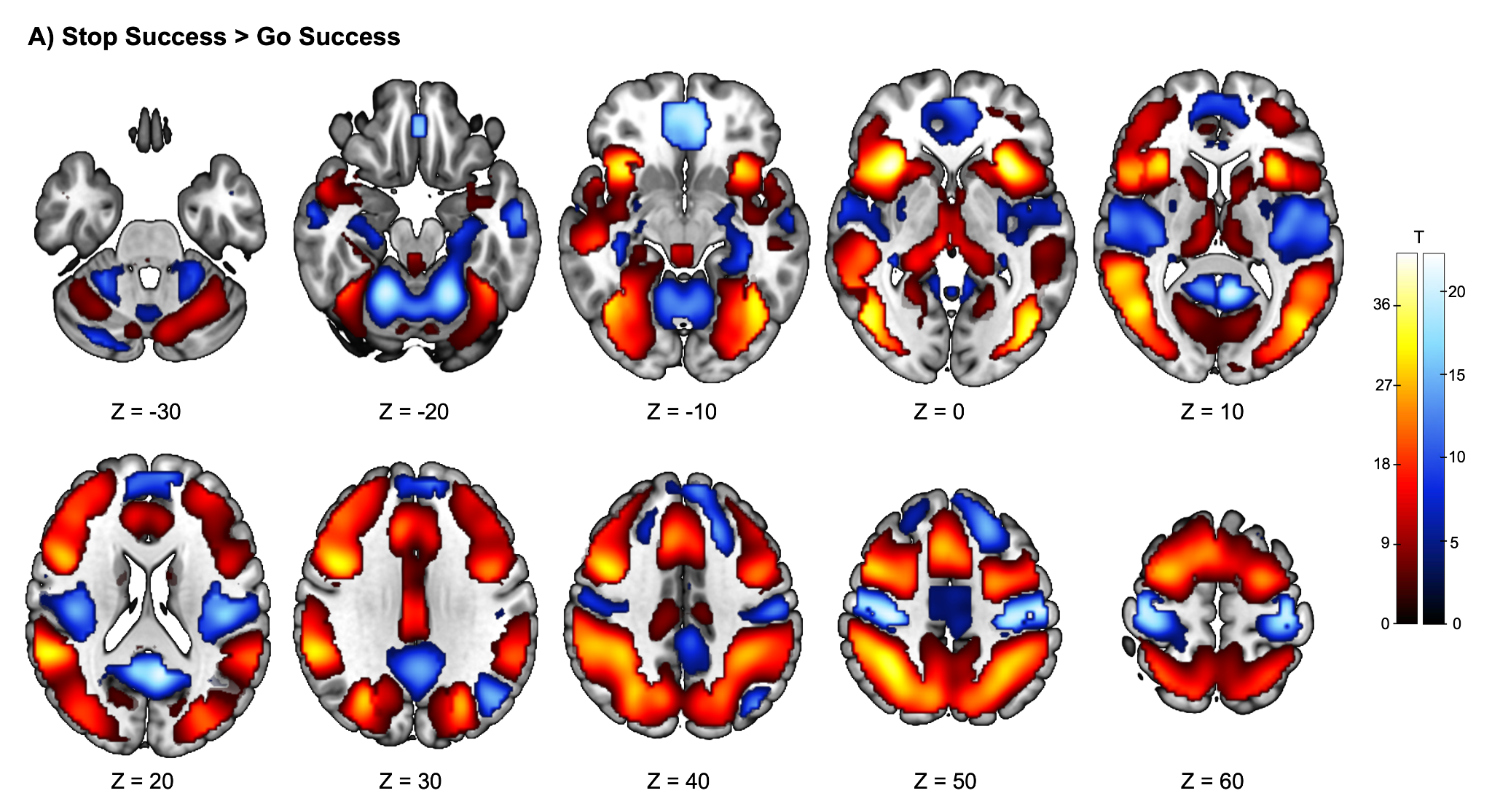

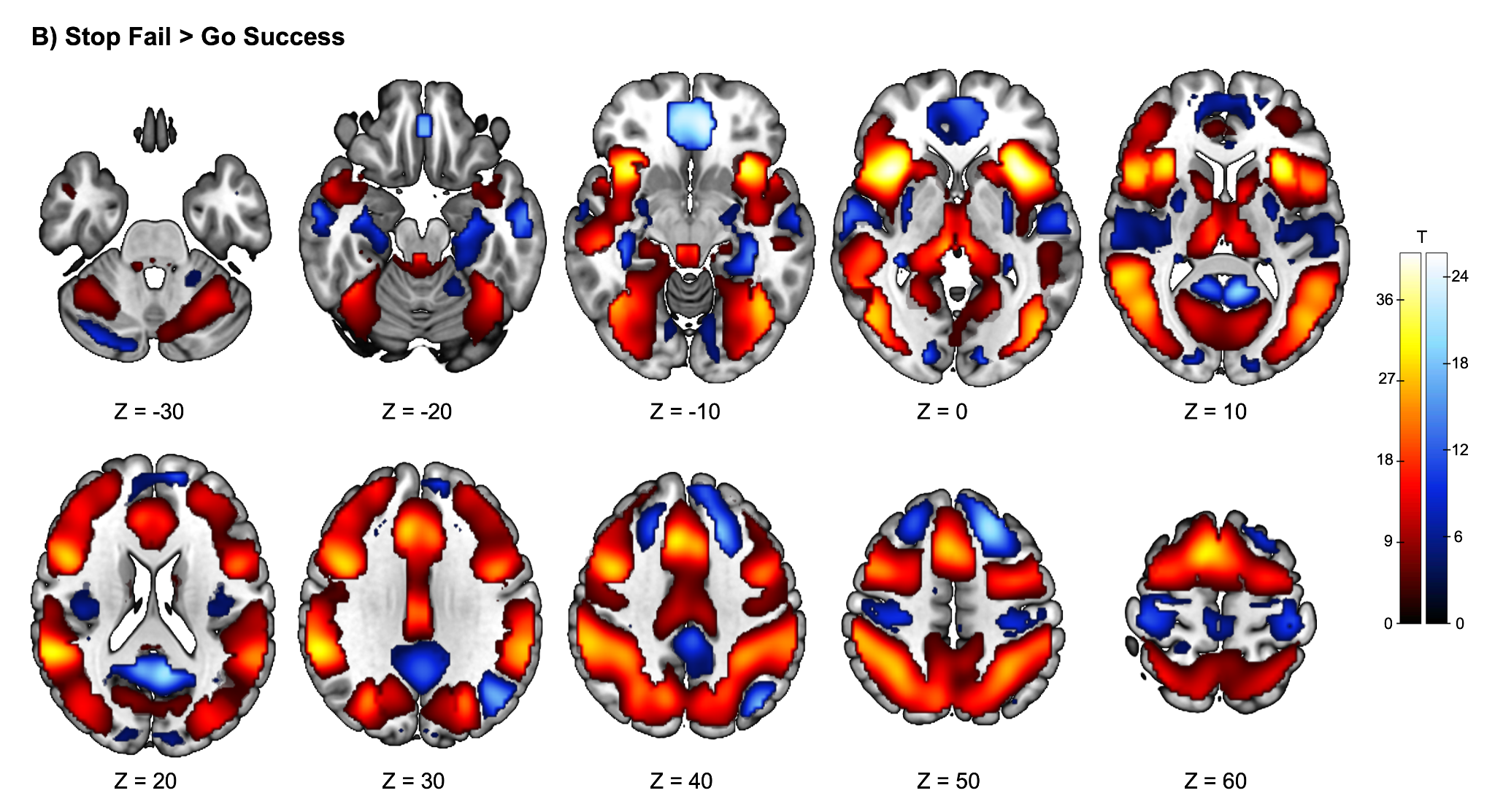

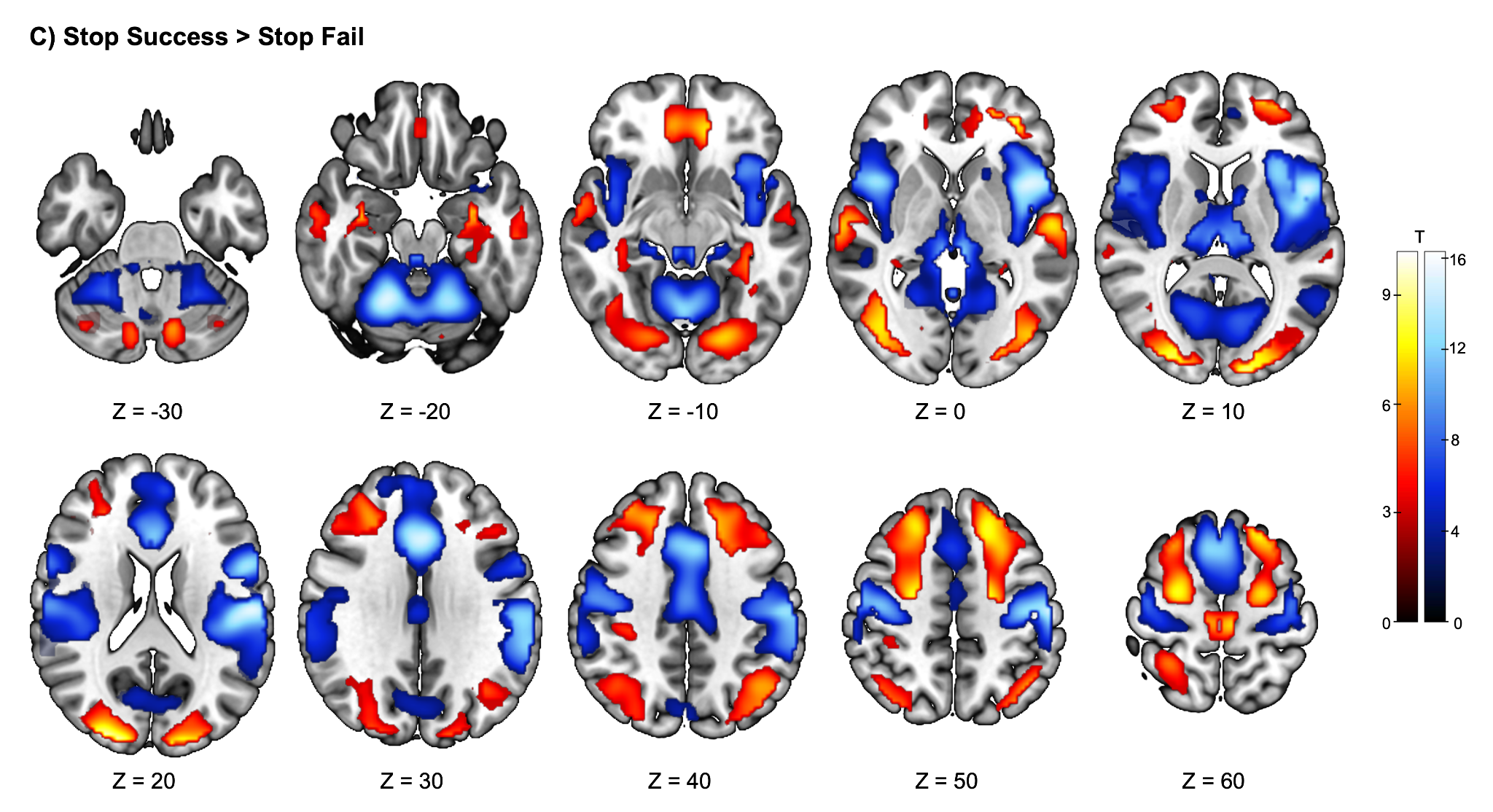

**Figure S2**. Task-elicited activation based on one-sample t-tests (*n* = 589) from contrasts **A)** Successful Stop > Successful Go; **B)** Failed Stop > Successful Go; and **C)** Successful Stop > Failed Stop. Significant activations were determined by a cluster-defining threshold of *p* < 0.001 (uncorrected), corrected to a cluster-extent threshold of *p_FWE_* < 0.05.

**Table S1**. Variance inflation factors (VIFs) for all predictors included in the linear regression models, calculated to assess potential multicollinearity.

| **Variable** | **Variance inflation factor (VIF)** |
| --- | --- |
| DSM-5: Impaired Control | 1.62 |
| DSM-5: Social Impairments | 1.26 |
| DSM-5: Risky Use | 1.26 |
| DSM-5: Physical Dependence | 1.16 |
| Age | 1.24 |
| Sex | 1.02 |
| Site | 1.07 |

Note: Variance inflation factors (VIFs) did not indicate problematic multicollinearity based on commonly used thresholds (O’Brien, 2007).

**Table S2**. Associations between behavioral inhibitory control performance parameters in Stop-Signal Task (SST) and AUD symptom domains in DSM-5 (n = 589).

| **SST performance measure** | **Impaired Control** | | | | | **Social Problems** | | | | | **Risky Use** | | | | | **Physical Dependence** | | | | |
| --- | --- | --- | --- | --- | --- | --- | --- | --- | --- | --- | --- | --- | --- | --- | --- | --- | --- | --- | --- | --- |
|  | ***β*** | **95% CI** | **b (SE)** | ***p (p_FDR_)*** | ***f^2^*** | ***β*** | **95% CI** | **b(SE)** | ***p (p_FDR_)*** | ***f^2^*** | ***β*** | **95% CI** | **b(SE)** | ***p (p_FDR_)*** | ***f^2^*** | ***β*** | **95% CI** | **b(SE)** | ***p (p_FDR_)*** | ***f^2^*** |
| Stop Signal reaction time (SSRT) | 0.00003 | [-0.10; 0.10] | -0.001  (2.61) | .999  (.999) | <0.001 | 0.0008 | [-0.09; 0.09] | 0.07  (4.05) | .985 (.985) | <0.001 | 0.05 | [-0.04; 0.13] | 4.02  (3.86) | .299 (.448) | 0.002 | 0.04 | [-0.04; 0.13] | 6.93  (6.69) | .301 (.534) | 0.002 |
| Balanced integration score (BIS) | -0.12 | [-0.22; -0.03] | -0.14  (0.06) | **.015 (.026)** | 0.01 | -0.007 | [-0.09; 0.08] | -0.01  (0.09) | .870 (.985) | <0.001 | -0.07 | [-0.15; 0.02] | -0.13  (0.08) | .121 (.364) | 0.004 | 0.03 | [-0.06; 0.11] | 0.09  (0.15) | .534 (.534) | 0.001 |
| Post-error slowing (PES) | -0.12 | [-0.23; -0.02] | -4.63  (1.95) | **.018 (.026)** | 0.01 | 0.07 | [-0.02; 0.16] | 4.32  (3.02) | .153 (.458) | 0.004 | -0.03 | [-0.12; 0.06] | -1.96  (2.88) | .497 (.497) | 0.001 | 0.04 | [-0.05; 0.12] | 4.16  (4.98) | .404 (.534) | 0.001 |

Note: Separate models were computed for each SST performance measure, incorporating all DSM-5 symptom domains (impaired control, social problems, risky use, physical dependence) as well as age, sex, and site as covariates. *p*-values were FDR-corrected for the number of SST measures. Bold printed *p*-values indicate significant associations after multiple testing corrections.

**Table S3**. Whole-brain associations between social problems symptom severity and neural activation during successful response inhibition (SS > SGo), controlling for other AUD symptom domains, site, and mean FD.

| **Contrast** | **Cluster No.** | **Peak region** | **Hemisphere** | **MNI** | **k_E_** | **T** | **Z** | ***p_FWE_*** |
| --- | --- | --- | --- | --- | --- | --- | --- | --- |
| SS > SGo | 1 | Occipital_Mid | R | 37, -81, 28 | 506 | 4.98 | 4.93 | .001 |

Note: Significant activation was determined by a cluster-defining threshold of *p* < 0.001 (uncorrected), corrected to a cluster-extent threshold of *p_FWE_* < 0.05. *k_E_* = number of voxels in cluster; T and Z are peak-level. Region labels are based on AAL3.

**Table S4**. Associations between alcohol drinking behaviors assessed at Baseline (BL) and AUD symptom domains in DSM-5 (n = 589).

| **Drinking outcome** | **Impaired Control** | | | | | **Social Problems** | | | | | **Risky Use** | | | | | **Physical Dependence** | | | | |
| --- | --- | --- | --- | --- | --- | --- | --- | --- | --- | --- | --- | --- | --- | --- | --- | --- | --- | --- | --- | --- |
|  | ***β*** | **95% CI** | **b(SE)** | ***p (p_FDR_)*** | ***f^2^*** | ***β*** | **95%  CI** | **b(SE)** | ***p (p_FDR_)*** | ***f^2^*** | ***β*** | **95% CI** | **b(SE)** | ***p (p_FDR_)*** | ***f^2^*** | ***β*** | **95% CI** | **b(SE)** | ***p (p_FDR_)*** | ***f^2^*** |
| BL: Average amount of daily alcohol intake | 0.20 | [0.10; 0.29] | 6.67  (1.69) | **< .001**  **(< .001)** | 0.027 | 0.11 | [0.02; 0.19] | 6.47  (2.62) | .**014 (.042)** | 0.010 | 0.14 | [0.05; 0.22] | 7.85  (2.50) | **.002 (.003)** | 0.017 | 0.05 | [-0.03; 0.13] | 4.90  (4.33) | .258 (.574) | 0.002 |
| BL: Frequency of alcohol drinking | 0.29 | [0.20; 0.38] | 6.60  (1.06) | **< .001 (< .001)** | 0.066 | 0.05 | [-0.04; 0.13] | 1.81  (1.65) | .273 (.273) | 0.002 | 0.22 | [0.14; 0.31] | 8.56  (1.57) | **< .001 (< .001)** | 0.051 | 0.02 | [-0.06; 0.10] | 1.53  (2.74) | .574 (.574) | 0.001 |
| BL: Frequency of binge-drinking | 0.20 | [0.11; 0.30] | 3.35  (0.83) | **< .001 (< .001)** | 0.028 | 0.07 | [-0.01; 0.16] | 2.08  (1.28) | .104 (.156) | 0.005 | 0.13 | [0.05; 0.22] | 3.68  (1.22) | **.003 (.003)** | 0.016 | -0.02 | [-0.11; 0.06] | -1.23  (2.11) | .561 (.574) | 0.001 |

Note: Separate models were computed for each drinking outcome, incorporating all DSM-5 symptom domains (impaired control, social problems, risky use, physical dependence) as well as age, sex, and site as covariates. *p*-values were FDR-corrected for the number of drinking outcomes. Bold printed *p*-values indicate significant associations after multiple testing corrections.

**Table S5**. Associations between alcohol drinking behaviors assessed during 12-months Ecological Momentary Assessment (EMA) and AUD symptom domains in DSM-5 (n = 389).

| **Drinking outcome** | **Impaired Control** | | | | | **Social Problems** | | | | | **Risky Use** | | | | | **Physical Dependence** | | | | |
| --- | --- | --- | --- | --- | --- | --- | --- | --- | --- | --- | --- | --- | --- | --- | --- | --- | --- | --- | --- | --- |
|  | ***Effect size*** | **95% CI** | **b(SE)** | ***p (p_FDR_)*** | ***f^2^*** | ***Effect size*** | **95% CI** | **b(SE)** | ***p (p_FDR_)*** | ***f^2^*** | ***Effect size*** | **95% CI** | **b(SE)** | ***p (p_FDR_)*** | ***f^2^*** | ***Effect size*** | **95% CI** | **b(SE)** | ***p (p_FDR_)*** | ***f^2^*** |
| EMA: Daily amount of alcohol drinking | *β =* 0.08 | [0.02; 0.13] | 3.82  (1.32) | **.004**  **(.012)** | 0.006 | *β = -*0.001 | [-0.05; 0.05] | -0.04  (1.57) | .979 (.979) | <0.001 | *β =* 0.03 | [-0.02; 0.08] | 2.00  (1.68) | .233 (.497) | 0.001 | *β =* 0.10 | [0.05; 0.15] | 9.99 (2.64) | **< .001 (.001)** | 0.010 |
| EMA: Frequency of alcohol drinking | OR = 1.03 | [0.88; 1.20] | 0.03  (0.08) | .733 (.733) | <0.001 | OR = 0.90 | [0.75; 1.08] | -0.10  (0.09) | .271 (.814) | 0.002 | OR = 0.99 | [0.81; 1.21] | -0.01  (0.10) | .928 (.928) | <0.001 | OR = 1.53 | [1.13; 2.09] | 0.43  (0.16) | **.007 (.007)** | 0.008 |
| EMA: Frequency of binge-drinking | OR = 1.22 | [1.04; 1.43] | 0.20  (0.08) | **.014 (.021)** | 0.006 | OR = 0.95 | [0.79; 1.15] | -0.05  (0.10) | .616 (.924) | <0.001 | OR = 1.10 | [0.90; 1.35] | 0.10  (0.10) | .331 (.497) | 0.001 | OR = 1.70 | [1.24; 2.34] | 0.53  (0.16) | **.001 (.001)** | 0.012 |

Note: Separate models were computed for each drinking outcome, incorporating all DSM-5 symptom domains (impaired control, social problems, risky use, physical dependence) as well as age, sex, and site as covariates. *p*-values were FDR-corrected for the number of drinking outcomes. Bold printed *p*-values indicate significant associations after multiple testing corrections.

**Table S6**. Exploration of associations between neural activation in clusters associated with impaired control symptoms during inhibitory control demands and drinking behavior.

| **Drinking outcome** | ***Successful Stop > Successful Go*** | | | | | | | | | | | | | | | ***Successful Stop >  Failed Stop*** | | |
| --- | --- | --- | --- | --- | --- | --- | --- | --- | --- | --- | --- | --- | --- | --- | --- | --- | --- | --- |
|  | ***Cluster 1*** | | | ***Cluster 2*** | | | ***Cluster 3*** | | | ***Cluster 4*** | | | ***Cluster 5*** | | | ***Cluster 1*** | | |
|  | ***Effect size*** | **95% CI** | ***p (p_FDR_)*** | ***Effect size*** | **95% CI** | ***p (p_FDR_)*** | ***Effect size*** | **95% CI** | ***p (p_FDR_)*** | ***Effect size*** | **95% CI** | ***p (p_FDR_)*** | ***Effect size*** | **95% CI** | ***p (p_FDR_)*** | ***Effect size*** | **95% CI** | ***p (p_FDR_)*** |
| BL: Average amount of daily alcohol intake | *β =* 0.02 | [-0.06; 0.10] | .645  (.680) | *β =* 0.03 | [-0.05; 0.11] | .434 (.695) | *β =* 0.03 | [-0.06; 0.11] | .532 (.623) | *β =* 0.05 | [-0.03; 0.13] | .206 (.478) | *β =* 0.03 | [-0.05; 0.11] | .420 (.840) | *β =* 0.03 | [-0.05; 0.11] | .516 (.946) |
| BL: Frequency of alcohol drinking | *β =* 0.07 | [-0.01; 0.15] | .091 (.546) | *β =* 0.06 | [-0.02; 0.15] | .118 (.695) | *β =* 0.05 | [-0.03; 0.13] | .261  (.623) | *β =* 0.09 | [0.01; 0.17] | .023 (.138) | *β =* 0.03 | [-0.05; 0.12] | .404 (.840) | *β =* 0.03 | [-0.05; 0.11] | .435 (.946) |
| BL: Frequency of binge-  drinking | *β =* 0.03 | [-0.05; 0.11] | .442 (.680) | *β =* 0.03 | [-0.05; 0.11] | .463 (.695) | *β =* 0.02 | [-0.06; 0.10] | .617  (.623) | *β =* 0.05 | [-0.03; 0.13] | .239 (.478) | *β =* 0.01 | [-0.07; 0.09] | .796 (.882) | *β =* 0.00 | [-0.08; 0.08] | .964 (.964) |
| EMA: Daily amount of alcohol drinking | *β = -*0.02 | [-0.07; 0.03] | .488 (.680) | *β = -*0.00 | [-0.06; 0.05] | .854 (.854) | *β = -*0.01 | [-0.06; 0.04] | .623  (.623) | *β = -*0.00 | [-0.06; 0.04] | .739 (.739) | *β =* 0.00 | [-0.04; 0.06] | .821 (.882) | *β = -*0.00 | [-0.06; 0.04] | .788 (.946) |
| EMA: Frequency of alcohol drinking | OR = 1.08 | [0.91; 1.29] | .391 (.680) | OR = 1.12 | [0.91; 1.37] | .274 (.695) | OR = 1.05 | [0.87; 1.27] | .595  (.623) | OR = 1.07 | [0.89; 1.28] | .466 (.677) | OR *=* 1.10 | [0.91; 1.34] | .314 (.840) | OR *=* 1.03 | [0.87; 1.22] | .740 (.946) |
| EMA: Frequency of binge- drinking | OR = 0.96 | [0.81; 1.14] | .680 (.680) | OR = 0.97 | [0.80; 1.18] | .754 (.854) | OR = 0.95 | [0.79; 1.14] | .551  (.623) | OR = 0.95 | [0.79; 1.13] | .564 (.677) | OR *=* 1.01 | [0.84; 1.22] | .882 (.882) | OR *=* 0.97 | [0.82; 1.15] | .757 (.946) |

Note: Separate models were computed for each drinking outcome and each significant brain activation cluster. *p*-values were FDR-corrected for the number of drinking outcomes.
